# Multi-View Integrative Approach For Imputing Short-Chain Fatty Acids and Identifying Key factors predicting Blood SCFA

**DOI:** 10.1101/2024.09.25.614767

**Authors:** Anqi Liu, Bo Tian, Chuan Qiu, Kuan-Jui Su, Lindong Jiang, Chen Zhao, Meng Song, Yong Liu, Gang Qu, Ziyu Zhou, Xiao Zhang, Shashank Sajjan Mungasavalli Gnanesh, Vivek Thumbigere-Math, Zhe Luo, Qing Tian, Li-Shu Zhang, Chong Wu, Zhengming Ding, Hui Shen, Hong-Wen Deng

## Abstract

Short-chain fatty acids (SCFAs) are the main metabolites produced by bacterial fermentation of dietary fiber within gastrointestinal tract. SCFAs produced by gut microbiotas (GMs) are absorbed by host, reach bloodstream, and are distributed to different organs, thus influencing host physiology. However, due to the limited budget or the poor sensitivity of instruments, most studies on GMs have incomplete blood SCFA data, limiting our understanding of the metabolic processes within the host. To address this gap, we developed an innovative multi-task multi-view integrative approach (M^2^AE, Multi-task Multi-View Attentive Encoders), to impute blood SCFA levels using gut metagenomic sequencing (MGS) data, while taking into account the intricate interplay among the gut microbiome, dietary features, and host characteristics, as well as the nuanced nature of SCFA dynamics within the body. Here, each view represents a distinct type of data input (i.e., gut microbiome compositions, dietary features, or host characteristics). Our method jointly explores both view-specific representations and cross-view correlations for effective predictions of SCFAs. We applied M^2^AE to two in-house datasets, which both include MGS and blood SCFAs profiles, host characteristics, and dietary features from 964 subjects and 171 subjects, respectively. Results from both of two datasets demonstrated that M^2^AE outperforms traditional regression-based and neural-network based approaches in imputing blood SCFAs. Furthermore, a series of gut bacterial species (e.g., *Bacteroides thetaiotaomicron* and *Clostridium asparagiforme*), host characteristics (e.g., race, gender), as well as dietary features (e.g., intake of fruits, pickles) were shown to contribute greatly to imputation of blood SCFAs. These findings demonstrated that GMs, dietary features and host characteristics might contribute to the complex biological processes involved in blood SCFA productions. These might pave the way for a deeper and more nuanced comprehension of how these factors impact human health.

## Introduction

Short-chain fatty acids (SCFAs) are vital metabolites produced by the bacterial fermentation of dietary fiber in the gastrointestinal tract ^1^. In a healthy gut, common bacteria such as Prevotella, Bacteroides, Ruminococcaceae, and Lachnospiraceae ^2, 3^ generate principal SCFAs, including acetate, propionate, and butyrate. These SCFAs are absorbed and distributed to various organs, influencing host physiology by maintaining an intestinal anaerobic environment and regulating energy metabolism ^4–7^. Given the effects of blood SCFAs on host health, understanding the factors that regulate their production is important for the development of strategies to modulate blood SCFA levels to promote health and prevent diseases ^8^.

Recent studies have shown that fermentative bacteria species such as *Faecalibacterium prausnitzii* and *Eubacterium rectale*, which are abundant in the human gut, could efficiently metabolize complex carbohydrates, particularly resistant starches, into butyrate ^9^. Wang *et al.* found that an imbalance of “Good” and “Bad” gut microbiota led to the attenuation of the bacterial metabolite SCFAs ^10, 11^. These findings demonstrated that gut microbiotas (GMs) are crucial determinants for blood SCFAs. Dietary features, specifically fibers and macronutrients (i.e., fat, protein, and carbohydrate) intake, are pivotal, as they determine the substrate availability for microbial fermentation in the gut, which subsequently impacts the synthesis of SCFAs ^12^. Meanwhile, host characteristics (e.g., age, race) can significantly modulate blood levels of SCFAs by influencing the synthesis, uptake, and utilization of blood SCFAs within the body, possibly through the direct or indirect regulation of metabolic processes and immune responses ^13^. For instance, the increase in blood SCFAs may be due to increased uptake of SCFAs in the colon, in part due to increased nutrient intake, a complete bypass of SCFA transporters and increased passive uptake of SCFAs ^14^. Despite the growing interest in SCFAs research, the majority of existing research does not fully account for the complex interplay among host characteristics, dietary features, and GMs, which are crucial for generating accurate results across a wide range of applications ^15^.

Due to the limited budget or the poor sensitivity of instruments, most of current studies focusing on the gut microbiome are lacking in complete measurements of blood SCFAs ^16^, which can limit subsequent analyses and conceivably results in the neglect of pivotal insights into metabolic processes within the host. Gut microbiome as measured by metagenomic sequencing (MGS) data can determine SCFA concentrations, influencing host phenotypes by affecting metabolism, immune responses, and energy homeostasis ^17, 18^. Integrating SCFA data with MGS data enables multi-omic analyses that reveal broader metabolic impacts and potential links between gut microbiota composition and physiological or disease-related outcomes ^19^. Hence, developing a model to impute blood SCFA levels using the MGS data is beneficial and essential for advancing our understanding in this field. To the best of our knowledge, the imputation of blood SCFAs in the host using the MGS data remains largely untapped, marking a significant gap in our comprehension of the dynamics and implications of blood SCFAs production. By developing predictive models that integrate gut microbial compositions with host characteristics and dietary features, we can better understand the complex interplay between these variables and their impacts on blood SCFA production, potentially paving the way for personalized interventions to optimize blood SCFA levels and promote overall well-being ^20^.

In this study, leveraging metagenomic sequencing technology and deep learning methods, we unveiled an innovative approach that captures the intricate interplays among GMs, dietary features, and host characteristics to impute the absolute abundances of human blood SCFAs. Our method addresses the challenge of incomplete SCFA data by imputing it using MGS data, which will further facilitate the integration of SCFAs and MGS. This integration will enable more comprehensive multi-omic analyses, providing deeper insights into the influence of gut microbial composition on SCFA levels and their subsequent impact on host phenotypes in future research. By applying our approach to two in-house generated datasets, we demonstrated our model outperforms traditional regression-based and neural-network based approaches in imputing blood SCFAs. Accurate imputation of incomplete blood SCFA data will enable researchers to conduct more comprehensive studies exploring metabolic processes and their potential implications for health.

## Methods

### Subject recruitment and sample collection

A total of 964 unrelated males, aged 20-51 years, were recruited for this study as the first dataset (Dataset 1). An additional 171 unrelated subjects, aged 20-85 years, were recruited for this study, forming the second dataset (Dataset 2). All the subjects were living in New Orleans, Louisiana and its surrounding areas. We excluded subjects who had chronic or recent temporary conditions (e.g., gastroenteritis or inter-continental travel in the past 3 months) that may have significantly disturbed gut microbiota compositions, as described previously ^21–27^. Each subject provided stool and blood samples for metagenomic and SCFA profiling, respectively. We used the OMNIgene•GUT (OMR-200) all-in-one system (DNA GenoTEK, Ottawa, CA) for stool sample collection. Stool samples were frozen at -80°C after sample procurement until DNA extraction. Serum (for 964 subjects) or plasma (for 171 subjects) was extracted from 10 ml of blood samples from each subject according to the protein precipitation protocol ^28^ developed for metabolomics analysis, aliquoted, and stored at -80°C until used for further analysis. The 964 subjects in the first dataset (Dataset 1) also completed three questionnaires—the Louisiana Osteoporosis Questionnaire, the Metagenomic Study Supplementary Questionnaire, and the Food Frequency Questionnaire—to provide relevant covariate information (e.g., demographic factors, lifestyle factors and dietary features). The 171 subjects in the second dataset (Dataset 2) completed only two questionnaires—the Louisiana Osteoporosis Questionnaire and the Metagenomic Study Supplementary Questionnaire—to provide similar covariate information (e.g., demographic factors, but part of lifestyle factors and dietary features). Each subject signed an informed consent, and the study protocols were approved by the Institutional Review Boards (IRBs) of Tulane University. All data were treated with confidentiality, ensuring the anonymity of the participants.

### Metagenome profiling

Metagenomic DNA was extracted from stool samples using the Nucleospin Soil kit (MACHEREY-NAGEL) according to manufacturer’s instructions, as previously described ^19, 29–33^. After a few washes, DNA was eluted with 50 μl elution buffer and stored at -80°C until used for further sequencing. For Dataset 1, 530 samples were sequenced at LC Sciences (Houston, TX), and 434 samples were sequenced at BGI Americas (Cambridge, MA). For Dataset 2, all the 171 subjects were sequenced at LC Sciences.

For the samples sequenced in LC Sciences (Houston, TX), the DNA library was constructed by TruSeq Nano DNA LT Library Preparation Kit (Illumina Inc.). And then we performed the paired-end 2×150 bp sequencing on an Illumina Hiseq 4000 platform at the LC Sciences following the vendor’s recommended protocol.

Raw sequencing reads were processed to obtain valid reads for further analysis. First, sequencing adapters were removed from sequencing reads using cutadapt v1.9 ^34^. Secondly, low quality reads were trimmed by fqtrim v0.94 ^35^ using a sliding-window algorithm. Thirdly, reads were aligned to the host genome using bowtie2 v2.2.0 ^36^ to remove host contamination. Once quality-filtered reads were obtained, they were *de novo* assembled to construct the metagenome for each sample by IDBA-UD v1.1.1 ^37^. All coding regions (CDS) of metagenomic contigs were predicted by MetaGeneMark v3.26 ^38^. CDS sequences of all samples were clustered by CD-HIT v4.6.1 ^39^ to obtain unigenes. Unigene abundance for a certain sample were estimated by TPM based on the number of aligned reads by bowtie2 v2.2.0 ^36^. The lowest common ancestor taxonomy of unigenes were obtained by aligning them against the NCBI NR database by DIAMOND v 0.9.^40^. For samples sequenced in BGI Americas, the sequencing library was generated using MGI

Easy Universal DNA Library Prep Set Kit (MGI Inc.). The established library was sequenced on BGI DNBSEQ platform using the 100 bp pair-end sample preparation protocol. Quality control (QC) of raw reads was performed using Fastp ^41^ to filter low-quality reads. The high-quality reads were aligned to the host genome using bowtie2 ^36^ to remove human reads. The gene profiles were generated by aligning high-quality sequencing reads to the 9.9M integrated gene catalog (IGC) by using the Human Microbiome Project Unified Metabolic Analysis Network (HUMAnN2) ^42^ .

### Serum/Plasma SCFA profiling

Eight SCFAs (acetic acid, propionic acid, isobutyric acid, butyric acid, 2-methylbutyric acid, isovaleric acid, valeric acid and hexanoic acid) in serum/plasma samples were analyzed by Metabolon Inc. (Durham, NC) using LC-MS/MS, as previously described ^29–33^. The serum/plasma samples were spiked with stable labelled internal standards and were homogenized and subjected to protein precipitation with an organic solvent. After centrifugation, an aliquot of the supernatant was derivatized. The reaction mixture was injected onto an Agilent 1290/AB Sciex QTrap 5500 LC MS/MS system equipped with a C18 reversed phase UHPLC column. The mass spectrometer was operated in negative mode using electrospray ionization (ESI). The peak area of the individual analyte product ions was measured against the peak area of the product ions of the corresponding internal standards. Quantitation was performed using a weighted linear least squares regression analysis generated from fortified calibration standards prepared immediately prior to each run. LC-MS/MS raw data were collected and processed using SCIEX OS-MQ software v1.7. Three levels of QC samples were prepared by diluting with phosphate-buffered saline (PBS) and/or spiking with stock solutions to obtain the appropriate concentrations for each level (low-concentration QC, medium-concentration QC, and high-concentration QC). Sample analysis was carried out in a 96-well plate format containing two calibration curves to determine SCFA concentrations and six QC samples (per plate) to monitor assay performance. Accuracy was evaluated using the corresponding QC replicates in the sample runs. QCs met acceptance criteria at all levels for all analytes. QC acceptance criteria are at least 50% of QC samples at each concentration level per analyte should be within ±20.0% of the corresponding historical mean, and at least 2/3 of all QC samples per analyte should fall within ±20.0% of the corresponding historical mean.

While SCFA levels were measured in Datasets 1 and 2 using serum and plasma samples, respectively, SCFA concentrations tend to be highly consistent between serum and plasma samples ^43, 44^. This consistency arises because both serum and plasma SCFAs reflect similar metabolic states and distributions in the bloodstream ^44^. Thus, validating the model on these two datasets is appropriate, as both serum and plasma SCFA data provide reliable and comparable insights into SCFA dynamics.

### Data preprocessing

To remove noise and experimental artifacts in the data and better interpret the results, proper preprocessing for each view data is essential. For gut metagenomic sequencing data, we kept only the GM species that exist in all the subjects for data harmonization across the two data sets. For dietary and clinical data, we filtered out variables with missing rate > 20% and kept all the variables that, to our knowledge ^45–48^, could be pertinent to the study. Missing data for dietary and host characteristics data were imputed using multiple imputation through R package ’mice’ ^49^. Missing values in SCFAs were imputed with the minimum of values of all subjects for each SCFA (missing rate <1%). We randomly selected 75% of the samples as the training set and the remaining 25% of the samples in the dataset as the test set. To avoid potential bias, the training data and testing data have been processed for data normalization separately. We applied log normalization to each type of data, transforming the features to reduce skewness and bring the values into a comparable scale.

### Overview of Multi-task Multi-View Attentive Encoders (M^2^AE) model

M^2^AE is a framework for prediction tasks with multi-view data as input. Each view corresponds to a distinct category of data input, i.e., gut microbiome compositions, dietary features, or host characteristics. The workflow of M^2^AE is shown in Fig. 1 and can be summarized into two components. (1) View-specific representation learning via attentive encoders. For each view, an attentive encoder is designed in a symmetric auto-encoder fashion, where the encoder part is composited with one graph convolutional module and two fully-connected layers for view-specific representation learning. (2) Multi-view integration via the View Interactive Network (VIN). A cross-view interactive tensor is calculated using the latent representations from all the view-specific networks. A VIN is then trained with the cross-view discovery tensor to produce the final predictions. VIN can effectively learn the intra-view and inter-view correlations in the higher-level space for better prediction with multi-view data. M^2^AE is an end-to-end model, where both view-specific attentive encoders and VIN module are trained jointly. We describe each component in detail in the following sections.

**Fig 1.**
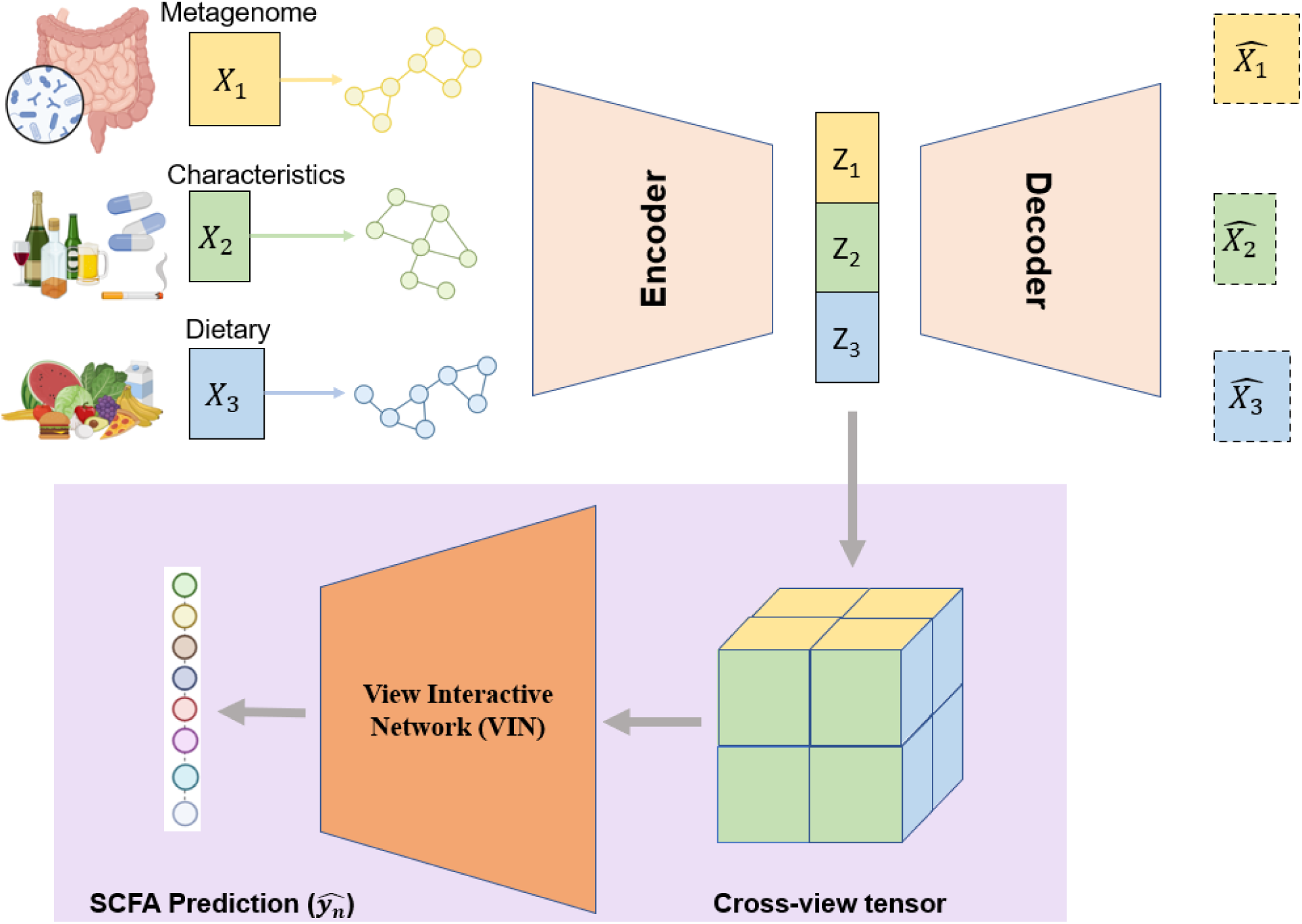
Overview of M^2^AE. M^2^AE is a framework for prediction tasks with multi-view data as input. The workflow of M^2^AE can be summarized into two components. (1) View-specific representation learning via attentive encoders. For each view, an attentive encoder was designed in a symmetric auto-encoder fashion, where the encoder part is composed with one graph convolutional module and two fully-connected layers for view-specific representation learning. (2) Multi-view integration via the View Interactive Network (VIN). A cross-view interactive tensor was calculated using the latent representations from all the view-specific networks. A VIN was then trained with the cross-view discovery tensor to produce the final predictions. VIN can effectively learn the intra-view and inter-view correlations in the higher-level space for better prediction with multi-view data. M^2^AE is an end-to-end model, where both view-specific attentive encoders and VIN module are trained jointly.

### Attentive encoders (AEs) for view-specific representation learning

We design each attentive encoder (AE) in an autoencoder manner with one encoder and one symmetric decoder. The encoder contains one graph convolutional module and two fully-connected layers for view-specific representation learning of each type of data input.

The graph convolutional module is implemented to map node features to low-dimensional space and utilizes a simple inner product layer to aggregate the features for feature embedding ^50^. By viewing each sample as a node, a view-specific graph can be constructed for each type of view by utilizing both the features (relative microbial abundance/dietary features/host characteristics) of each node and the relationships between nodes. Specifically, in each view, the input sample feature matrix **X** ∈ ℝ^*n*×*d*^contains the features of all samples, where *n* is the number of samples and *d* is the number of features. The input adjacency matrix **A** ∈ ℝ^*n*×*n*^characterizes the relationships between samples by computing the cosine similarity among pairs of nodes and edges. Thus, the graph convolutional module can be built by stacking multiple convolutional layers with each layer defined as:

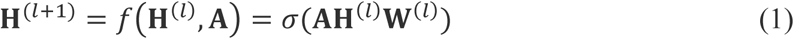

where **H**^(*l*)^ is the input of the *l*th layer and **W**^(*l*)^is the weight matrix of the *l*-th layer and **H**^(0)^ = **X**. *σ*(·) denotes a non-linear activation function. For **A**_*ij*_, it states the adjacency between node *i* and node *j* in the graph and is calculated as:

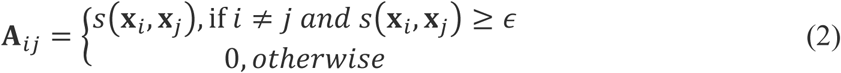

where **x***_i_* and **x***_j_* are the feature vectors of node *i* and node *j*, respectively. 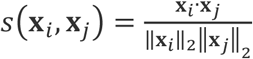 is the cosine similarity between node *i* and node *j*. The threshold *ɛ* is determined given a parameter *k*, which represents the average number of edges per node that are retained including self-connections:

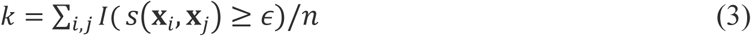

where *I*(·) is the indicator function. The parameter *k* in Eq. (3) is tuned over ^51^ {2, 5, 10} with the training data, and the same *k* value is adopted across all experiments on the same dataset. Note that for *k* = 1, **A** will turn out to be an identity matrix.

Our graph convolutional module will output a latent feature **F** per view, which is then fed into the subsequent two fully-connected layers generating a further latent feature **Z** for each view. Thus, the adjacency matrix in the decoder is calculated as **Â** = sigmoid(**ZZ**^**T**^) ^52^, which is sent to the decoder to reconstruct the original input.

For the model training, we aim to minimize the mean absolute error (MAE) between the input feature matrix **X** and the reconstructed matrix **X̂** for all views:

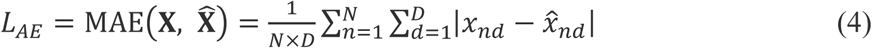

where MAE(·) represents the mean absolute error function. *x*_*nd*_ is the input feature of the *n*th sample and *d*th feature, *x̂*_*nd*_ is the predicted feature of the *n*th sample and *d*th feature.

So far, we have learned the view-specific representation **Z** and we will introduce to fuse each view for the final prediction task in the following section.

### VIN for multi-view integration

Current approaches leveraging multi-view data for biomedical prediction tasks traditionally either concatenate features from disparate views directly or fuse these features within a low-level feature space ^53–56^. However, properly aligning multiple views remains a consistent challenge, as improper alignment can have detrimental effects. On the other hand, view correlation discovery network (VCDN) ^57^ can exploit the higher-level cross-view correlations in class label level, as different types of data can provide unique distinctiveness for the production of SCFAs. Inspired by this, we develop VIN, which consolidates three latent features from gut microbiome, dietary features, and host characteristics to learn higher-level intra-view and cross-view correlations, thereby improving SCFA predictions.

For the latent representations of the *n*th sample from three types of views 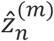, *m* = 1, 2, 3, we construct a cross-view interactive tensor *C*_*n*_, where each entry of *C*_*n*_ is calculated as:

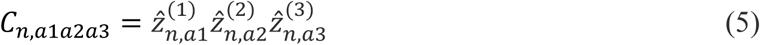

where 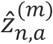 denotes the *a*th entry of 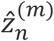. Then, the obtained tensor *C*_n_ is reshaped to a *c*^3^ dimensional vector **c**_*n*_ and is forwarded to the final prediction. VIN(·) is designed as a network with one graph convolutional layer and one fully-connected layer with the output dimension of *c* (In this case, we have eight SCFAs as outputs, so we set *c* = 8). We aim to minimize the mean absolute error between the predicted and ground-truth SCFAs as:

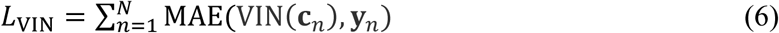

where **y**_*n*_ represents the absolute abundances of eight SCFAs in the *n*th sample, **N** represents the sample size.

To this end, VIN(·) could reveal the latent intra-view and cross-view correlations and help to improve the learning performance. By utilizing VIN(·) to integrate latent representations from different types of views, the final prediction made by M^2^AE is based on both the latent representation from each view and the learned cross-view correlation knowledge.

Overall, we optimize our M^2^AE by minimizing the attentive encoder loss and view interactive network losses in an iterative manner. During one epoch of the training process, we first fix VIN(·) and update *AE*_*m*_(·), *m* = 1, 2, 3, for each type of view to minimize the loss function 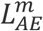 . Then we fix the view-specific AEs and update VIN(·) to minimize *L*_VIN_. View-specific AEs and VIN are updated alternately until convergence.

### Model performance evaluation

To evaluate the model’s performance in imputing blood SCFAs, we computed the mean absolute errors (MAE) and root mean squared errors (RMSE) for each subject. The average MAE and RMSE were then calculated by averaging these metrics across all subjects. We evaluated the models on five different randomly generated training and test splits, and the mean and standard deviation of the evaluation metrics across these five experiments were computed.

### Identification of influential factors for blood SCFAs

To identify significant factors for SCFAs, we defined a feature contribution score for each SCFA *p* across three different views as:

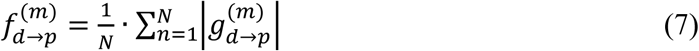

where 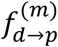 denotes the contribution score of the feature *d* to the SCFA *p* in view *m* and 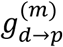 denotes the gradient of the SCFA *p* with respect to the input feature *d* in view *m*. Using this approach, we analyzed the contribution of each feature in different types of views on the test set. Features with the largest contribution scores in each view were considered to be the most important ones. Considering the inherent variability during training, we executed five repeated experiments in one dataset and reported the results by summing up the feature contribution scores across these five repeated experiments.

KEGG pathway analyses were conducted to identify significant biological pathways enriched in prominent bacterial species associated with SCFAs, by searching on the website of Kyoto Encyclopedia of Genes and Genomes (https://www.genome.jp/kegg/).

## Results

### Datasets

To validate the proposed M^2^AE model, we applied it to two different in-house datasets. We adopted the same data preprocessing pipeline described in the Methods section. For a fair comparison with existing approaches, we used the same methodology to construct the training and testing sets for evaluation.

**Dataset 1** consists of data from 964 unrelated males who provided both stool and blood samples for metagenomic and serum SCFAs profiling, along with dietary features, and host characteristics data. The basic characteristics of the samples are shown in Table 1. Features used to predict serum SCFAs include 194 gut bacterial species (relative abundance), 33 dietary features (e.g., intake of fruits, vegetables) and 17 host characteristics (e.g., age, race). The host characteristics and dietary habits used in Dataset 1 are listed in Supplementary Table 1.

**Table 1.**
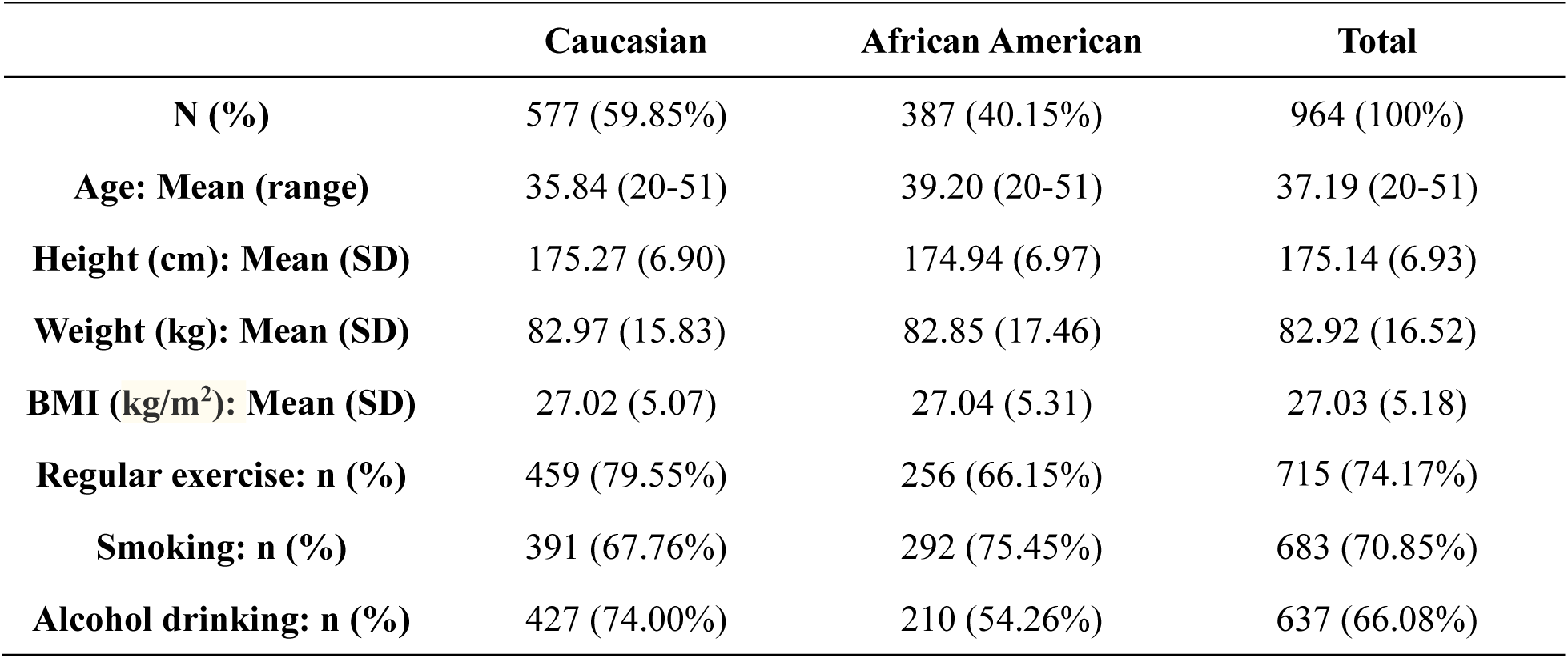
Sample Characteristics for Dataset 1 (964 subjects).

|  | Caucasian | African American | Total |
| --- | --- | --- | --- |
| **N (%)** | 577 (59.85%) | 387 (40.15%) | 964 (100%) |
| **Age: Mean (range)** | 35.84 (20-51) | 39.20 (20-51) | 37.19 (20-51) |
| **Height (cm): Mean (SD)** | 175.27 (6.90) | 174.94 (6.97) | 175.14 (6.93) |
| **Weight (kg): Mean (SD)** | 82.97 (15.83) | 82.85 (17.46) | 82.92 (16.52) |
| **BMI (kg/m<sup>2</sup>): Mean (SD)** | 27.02 (5.07) | 27.04 (5.31) | 27.03 (5.18) |
| **Regular exercise: n (%)** | 459 (79.55%) | 256 (66.15%) | 715 (74.17%) |
| **Smoking: n (%)** | 391 (67.76%) | 292 (75.45%) | 683 (70.85%) |
| **Alcohol drinking: n (%)** | 427 (74.00%) | 210 (54.26%) | 637 (66.08%) |

**Dataset 2** includes data from 171 unrelated subjects who provided both stool and blood samples for metagenomic profiling and plasma SCFAs profiling, along with dietary features and host characteristics data. The basic characteristics of these samples are presented in Table 2. Features used to predict plasma SCFAs include 646 gut bacterial species (relative abundance), 3 dietary features (e.g., intake of milk and yogurt), and 11 host characteristics (e.g., age, gender, and race). The host characteristics and dietary habits used in Dataset 2 are listed in Supplementary Table 2.

**Table 2.** Sample Characteristics for Dataset 2 (171 subjects).

|  | **Male (58 (33.92%))** |  | **Female (113 (66.08%))** |  |  |
| --- | --- | --- | --- | --- | --- |
|  | **Caucasian** | **African American** | **Caucasian** | **African American** | **Total** |
| **N (%)** | 19 (77.19%) | 39 (22.81%) | 77 (78.95%) | 36 (21.05%) | 171 (100%) |
| **Age: Mean (range)** | 58.16 (44-76) | 58.13 (51-72) | 41.66 (20-85) | 40.00 (21-69) | 46.90 (20-85) |
| **Height (cm): Mean (SD)** | 173.32 (8.07) | 172.93 (7.57) | 161.80 (6.96) | 165.52 (6.33) | 166.40 (8.63) |
| **Weight (kg): Mean (SD)** | 85.41 (14.12) | 83.72 (18.85) | 66.10 (14.53) | 77.34 (17.30) | 74.63 (17.97) |
| **BMI (kg/m<sup>2</sup>): Mean (SD)** | 28.53 (4.93) | 27.81 (4.90) | 25.24 (5.27) | 28.20 (6.02) | 26.82 (5.47) |
| **Regular exercise: n (%)** | 29 (83.04%) | 29 (16.96%) | 88 (85.38%) | 25 (14.62%) | 124 (72.51%) |
| **Smoking: n (%)** | 35 (86.55%) | 23 (13.45%) | 103 (94.15%) | 10 (5.85%) | 75 (43.86%) |
| **Alcohol drinking: n (%)** | 41 (90.06%) | 17 (9.94%) | 98 (91.23%) | 15 (8.77%) | 110 (64.33%) |

### M^2^AE outperformed existing multi-view integration prediction methods

As shown in Table 3, we compared the prediction performance of M^2^AE with the following existing regression algorithms for our data: (1) K-nearest neighbor regression (KNN), (2) Random forest regression (RF), (3) Gradient boosting-based regression (XGBoost), (4) Fully-connected neural network (NN) regression and (5) Linear regression. Deep fully-connected NN were also trained with MAE loss. Among the compared methods, KNN, RF, XGBoost, and NN were trained with the direct concatenation of the processed multi-view data as input. All methods were trained with the same processed data. The average MAE and average RMSE across all subjects were computed to compare the performance of different models. The choices for each hyper-parameter are relegated to Supplementary Table 3.

**Table 3.** Performance Comparison of different models for Dataset 1 and Dataset 2.

|  | Dataset 1 |  | Dataset 2 |  |
| --- | --- | --- | --- | --- |
| Methods | Mean MAE | Mean RMSE | Total MAE | Total RMSE |
| Linear Regression | $0.641 \pm 0.016$ | $0.813 \pm 0.020$ | $0.495 \pm 0.016$ | $0.620 \pm 0.002$ |
| RF | $0.481 \pm 0.012$ | $0.619 \pm 0.014$ | $0.393 \pm 0.018$ | $0.494 \pm 0.004$ |
| NN | $0.951 \pm 0.100$ | $1.170 \pm 0.121$ | $2.232 \pm 0.160$ | $2.673 \pm 0.062$ |
| KNN | $0.518 \pm 0.011$ | $0.662 \pm 0.014$ | $0.423 \pm 0.021$ | $0.530 \pm 0.005$ |
| XGBoost | $0.529 \pm 0.013$ | $0.679 \pm 0.013$ | $0.447 \pm 0.021$ | $0.569 \pm 0.005$ |
| AE(MLP) | $0.458 \pm 0.007$ | $0.589 \pm 0.010$ | $0.392 \pm 0.021$ | $0.502 \pm 0.004$ |
| AE(GCN) | $0.467 \pm 0.006$ | $0.601 \pm 0.008$ | $0.385 \pm 0.018$ | $0.494 \pm 0.004$ |
| AE(2GCN+1MLP) | $0.470 \pm 0.015$ | $0.602 \pm 0.015$ | $0.388 \pm 0.021$ | $0.498 \pm 0.003$ |
| VIN(MLP) | $0.458 \pm 0.011$ | $0.590 \pm 0.015$ | $0.384 \pm 0.019$ | $0.493 \pm 0.003$ |
| VIN(GCN) | $0.597 \pm 0.029$ | $0.731 \pm 0.032$ | $0.384 \pm 0.019$ | $0.493 \pm 0.003$ |
| M<sup>2</sup>AE | **$0.449 \pm 0.010$** | **$0.581 \pm 0.012$** | **$0.382 \pm 0.017$** | **$0.489 \pm 0.030$** |

Note: Bold represents the best performance in different criteria.

We observed that M^2^AE outperformed the other methods in the prediction tasks by showing the smallest mean MAE and mean RMSE in both Dataset 1 and Dataset 2 (Table 3), indicating the superior learning capability of M^2^AE. Interestingly, although deep learning-based methods have shown great promises in regression applications, the deep learning-based method NN did not show clear improvements over other approaches. This observation suggested that proper design of deep learning algorithms specific to multi-view integration applications was to some degree required to achieve superior prediction performance.

### M^2^AE outperformed its variations and other methods in SCFA prediction tasks

M^2^AE integrates view-specific learning via AEs with cross-view interactive fusion via VIN for effective SCFA predictions. To examine the necessity of AEs and VIN for effective SCFA predictions, we performed extensive ablation studies of our proposed method where two additional variations of M^2^AE were compared. (1) AutoEncoder_VIN: a) Fully-connected NNs with the same number of layers and the same dimensions of hidden layers as the encoder part in M^2^AE were used for view-specific representation learning; b) GCNs with the same number of layers and the same dimensions of hidden layers as the encoder part in M^2^AE were used for view-specific representation learning; c) An AE containing two graph convolutional layers and one fully-connected layer with the same dimensions of hidden layers as the encoder part in M^2^AE were used for view-specific representation learning. The multi-view integration component utilized VIN, which was the same as M^2^AE. (2) AE_NN/GCN: the view-specific representation component utilized one graph convolutional layer and two fully-connected layers, which was the same as M^2^AE. a) A fully-connected NN with the same number of layers and the same dimensions of hidden layers as VIN was used for multi-view integration; b) A GCN with the same number of layers and the same dimensions of hidden layers as VIN was used for multi-view integration. Note that Autoencoder_VIN itself is also a novel approach. To the best of our knowledge, there is no existing method that applies AEs to multi-view data integration and imputation problems.

As shown in Table 3, we observed that M^2^AE outperformed Autoencoder_VIN and AE_NN/GCN in all prediction tasks across both Datasets 1 and 2. The better performance of average MAE and average RMSE in M^2^AE than AE_NN/GCN indicates that our usage of VIN combines one graph convolutional layer and one fully-connected layer for multi-view integration and prediction tasks makes important contributions to the performance boost of M^2^AE comparing with existing methods. Compared with traditional NN and GCN that only learn from view information from one pathway, the AE further exploits the graph structural information within the data using a more flexible way. This can be essential to a more comprehensive understanding of the type of view as it captures the connections and correlations among samples. Therefore, AEs were needed for effective view-specific representation learning to fully exploit the advantages of VIN, and these two components could be trained jointly to achieve superior results for multi-view prediction tasks across both datasets.

### Performance of M^2^AE under different types of views

To further demonstrate the necessity of integrating multiple types of data to boost the prediction performance, we compared the prediction performance of M^2^AE with three types of views (MGS + host characteristics + dietary features), M^2^AE with two types of views (MGS + host characteristics, MGS + dietary features, and host characteristics + dietary features), and the view-specific AEs trained with single-view data before integration (MGS only, host characteristics only, and dietary features only). Figs. 2 and 3 show that by exploring the cross-view interactive fusion through VIN, the prediction performance was consistently improved by integrating prediction results from multiple views. Specifically, in all the prediction tasks, M^2^AE models trained with three types of views achieved the best performance compared with M^2^AE models trained with two types of views. Moreover, the M^2^AE models trained with two types of views both outperformed the single-view AE models.

**Fig 2.**
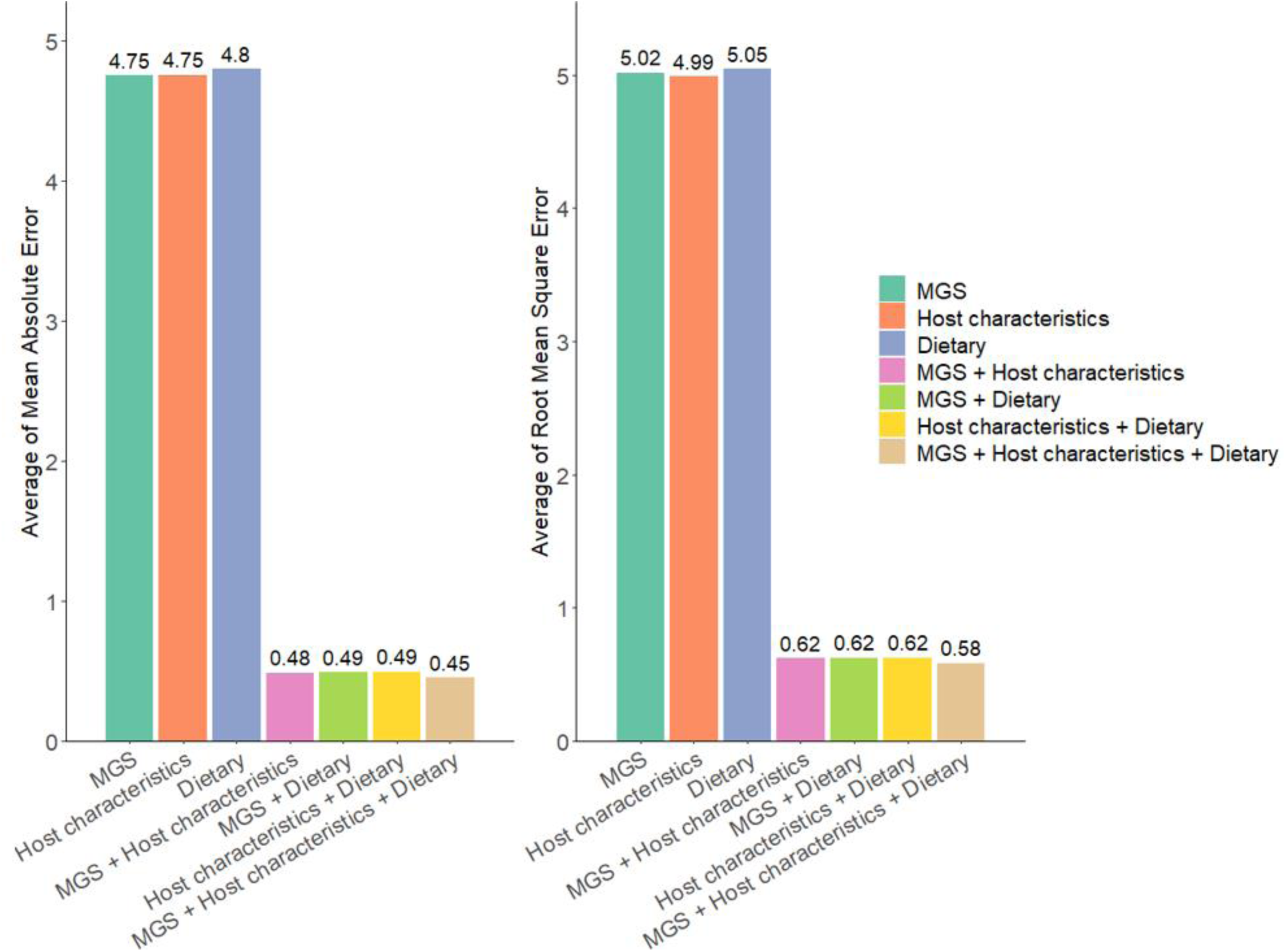
Performance comparison of multi-view data prediction via M^2^AE and single-view data prediction via Attentive Encoders in Dataset 1 (n = 5 experiments for each model).

**Fig 3.**
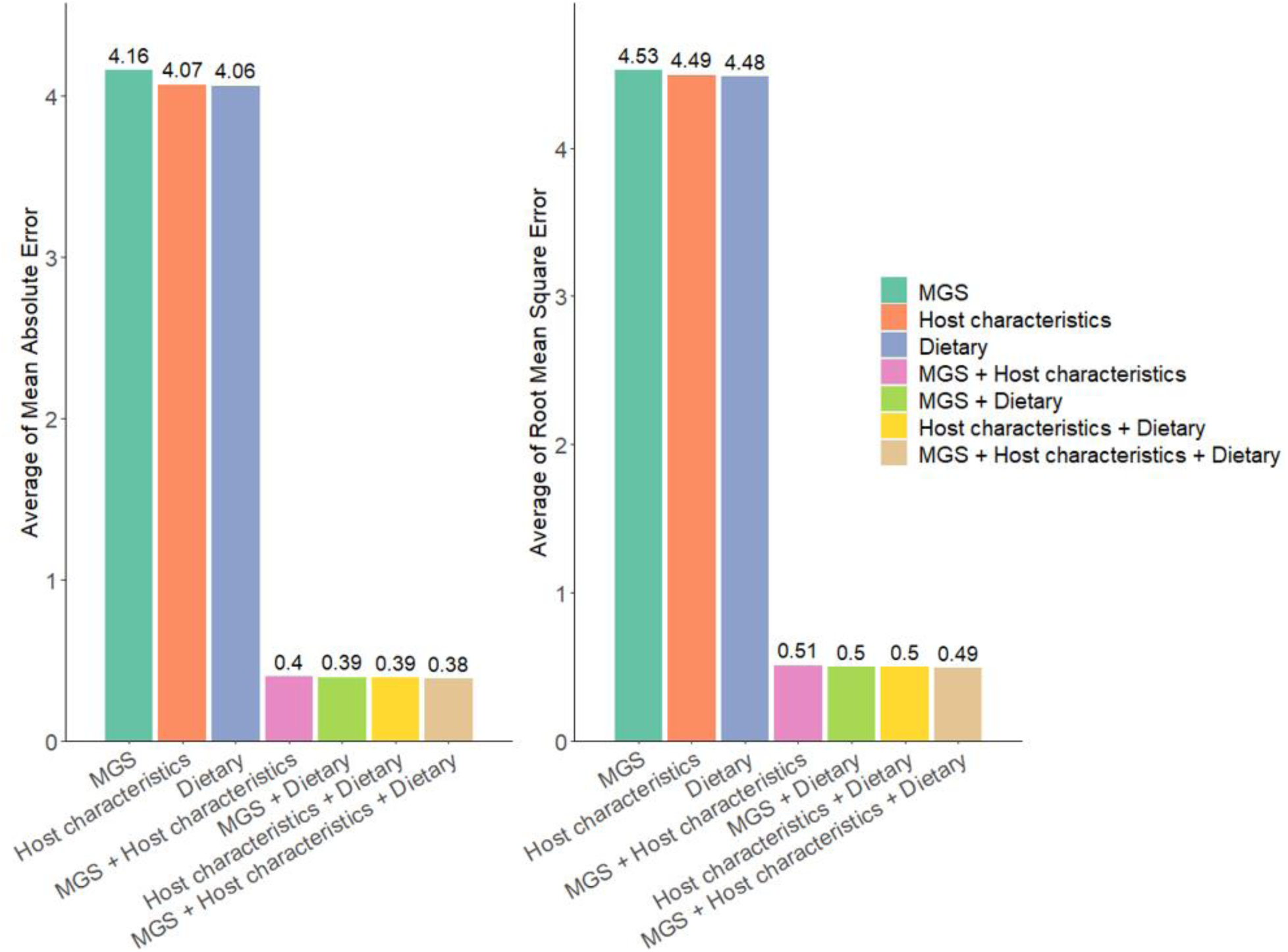
Performance comparison of multi-view data prediction via M^2^AE and single-view data prediction via Attentive Encoders in Dataset 2 (n = 5 experiments for each model).

It is well known that gut microbiome is closely influenced by host characteristics and dietary habits, as noted in previous studies ^12, 13^. This interdependence suggests that these factors, when considered together, may collectively enhance the predictive power of models. The results, therefore, strongly support the necessity of integrating multiple types of data in predictive models. By leveraging cross-view interactions and fusing diverse data types, we captured a more comprehensive understanding of the complex interplay between microbiome, host, and environmental factors, leading to significantly improved prediction accuracy. This multi-view approach is crucial for advancing personalized medicine and for a more profound understanding of complex biological phenomena.

Means of evaluation metrics from different experiments are shown in the figure. MGS + Host characteristics + Dietary refers to M^2^AE with three types of views combining MGS, host characteristics, and dietary features data. MGS + Host characteristics refers to M^2^AE with two types of views combining gut microbiomes and host characteristics. MGS + Dietary refers to M^2^AE with two types of views combining gut microbiomes and dietary features. Host characteristics + Dietary refers to M^2^AE with two types of views combining host characteristics and dietary features. MGS, host characteristics and dietary features refer to the view-specific attentive encoders trained with single-view gut microbiomes, host characteristics, and dietary features.

Means of evaluation metrics from different experiments are shown in the figure. MGS + Host characteristics + Dietary refers to M^2^AE with three types of views combining MGS, host characteristics, and dietary features data. MGS + Host characteristics refers to M^2^AE with two types of views combining gut microbiomes and host characteristics. MGS + Dietary refers to M^2^AE with two types of views combining gut microbiomes and dietary features. Host characteristics + Dietary refers to M^2^AE with two types of views combining host characteristics and dietary features. MGS, host characteristics and dietary features refer to the view-specific attentive encoders trained with single-view gut microbiomes, host characteristics, and dietary features.

### M^2^AE identified important factors associated with blood SCFAs

In our analysis, we identified key features influencing SCFA production by selecting top-ranked features from two datasets, as detailed in Supplementary Tables 4-11. Among these, *Faecalibacterium prausnitzii* and *Rothia mucilaginosa* emerged as significant contributors to various SCFAs production. Several species from the *Bacteroides* genus were also highlighted for their role in SCFA biosynthesis. For example, *Bacteroides thetaiotaomicron* and *Bacteroides fragilis* were major producers of acetic acid, while *Bacteroides vulgatus* was linked to valeric acid. *Bacteroides fragilis* was also associated with 2-methylbutyric acid and isobutyric acid, and *Bacteroides eggerthii* was found to be important for isovaleric acid production. Numerous species within the *Clostridium* genus were identified as key contributors to specific SCFAs. *Clostridium bolteae* was particularly relevant for isobutyric acid production, while *Clostridium asparagiforme* played a significant role in butyric acid synthesis. *Streptococcus salivarius* was associated with butyric acid production. Additionally, species like *Leuconostoc gelidum* was noted for its relevance to valeric acid biosynthesis pathways. KEGG pathway analysis further revealed that several of these bacterial species, including *Faecalibacterium prausnitzii*, *Rothia mucilaginosa*, *Bacteroides thetaiotaomicron*, *Bacteroides fragilis*, *Bacteroides vulgatus*, *Clostridium asparagiforme*, and *Leuconostoc gelidum*, are enriched in SCFA-related biological processes, such as fatty acid biosynthesis, degradation, and metabolism.

Moreover, host characteristics, such as gender, race, age, height, weight, physical activity levels, and the use of probiotics, antibiotics, and gastric acid-lowering medications, were found to correlate with SCFA levels (Supplementary Tables 4-11). Dietary habits, including the consumption of pickles, fruits, cereals, eggs, meat, fats, coffee, and chocolate, also significantly influenced SCFA production. Overall, the factors identified by the M^2^AE model showed substantial diversities between different SCFAs.

## Discussion

Recent development in high-throughput profiling technologies and integrative analysis of multi-view data offered advanced and powerful approaches to dissect complex biological problems. In this study, we pioneered an innovative approach, M^2^AE, for imputing the abundances of blood SCFAs, and performed multi-view prediction for blood SCFAs data by synthesizing the information of gut microbiome, dietary features and host characteristics. This method jointly explores view-specific representation and cross-view correlation for effective prediction, and demonstrated superior performance compared with other methods. M^2^AE also effectively identified prominent factors that showed strong associations with blood SCFAs.

Through literature mining, we found interesting evidence supporting the biological connections between these prominent factors and blood SCFAs and interesting relationships among some of these prominent factors.

### Gut microbiotas

Our analysis identified several GM species, such as *Bacteroides thetaiotaomicron, Bacteroides fragilis*, *Bacteroides vulgatus*, *Bacteroides eggerthii*, *Clostridium asparagiforme, Clostridium bolteae, Faecalibacterium prausnitzii, Rothia mucilaginosa, Streptococcus salivarius* and *Leuconostoc gelidum*, as significant contributors to the production of blood SCFAs. Specifically, *Bacteroides thetaiotaomicron* and *Bacteroides fragilis* are prominent contributors to acetic acid production due to their ability to ferment complex carbohydrates into intermediate metabolites, such as lactate and succinate, which can be further converted into SCFAs by other gut bacteria ^30, 58, 59^. *Bacteroides vulgatus* exhibited a negative association with blood valeric acid levels ^60^, possibly due to its negative interactions with other gut bacteria, which affect substrate availability for valeric acid production ^61^. *Bacteroides eggerthii* has been identified as a significant contributor to the production of isovaleric acid, primarily via leucine fermentation ^62^. Moreover, consistent with prior study, *Clostridium asparagiforme* plays a major role in butyrate production by fermenting glucose into lactate, which is then converted to butyrate by other bacteria ^63^. *Clostridium bolteae* can utilize valine through fermentation pathways, leading to the production of isobutyric acid as a metabolic byproduct ^64^. *Faecalibacterium prausnitzii* is positively correlated with butyric and valeric acid ^59, 65, 66^. *Rothia mucilaginosa* ferments glucose to produce acetate ^67^. As above, we identified a series of gut bacterial species that have been proved to play a role in SCFA production. In addition, we identified some novel putative factors that might affect blood SCFAs. For example, *Bacteroides fragilis* was associated with 2-methylbutyric acid and isobutyric acid, *Streptococcus salivarius* was associated with butyric acid production, and *Leuconostoc gelidum* was related to valeric acid. *Bacteroides* species contribute to amino acid metabolism ^68^, which might lead to the production of SCFAs, such as isobutyric acid and 2-methylbutyric acid. *Streptococcus salivarius*, primarily known for its presence in the oral cavity, can also inhabit the gut and metabolize carbohydrates via fermentation ^69^, potentially contributing to the production of butyric acid. Similarly, *Leuconostoc gelidum* is a lactic acid bacterium known for fermenting carbohydrates to produce lactic acid ^70^, which might serve as a substrate for other gut microbes, potentially leading to the production of valeric acid. These findings enhance our understanding of the complex interactions between GM and blood SCFA levels. Meanwhile, these insights could help in validating our model by supporting the observed associations between specific bacterial species and SCFA production, as well as their potential influence on systemic SCFA levels. However, more in-depth studies might be needed to further unravel the underlying mechanisms.

### Dietary features

Incorporating dietary features into our study enhanced our understanding in regulating the complex biological regulation of blood SCFAs production. Our findings demonstrated that the intake of various dietary components, such as pickles, fruits, cereals, eggs, meat, fat oil, coffee, and chocolate, influences blood SCFA levels. Diets could shape the microbiome by promoting the growth of bacteria that preferentially use the ingested nutrients ^71^. For instance, fermented foods like pickles and fiber-rich foods like fruits and cereals promote the growth of beneficial bacteria that ferment sugars and fibers into SCFAs, such as acetic, propionic, and butyric acids ^72, 73^. High- protein and high-fat diets, including eggs, steak, and fat oil, can promote the growth of *Bacteroides* species, which are adept at protein degradation and fat metabolism, thereby affecting SCFA production ^10, 74–76^. Additionally, coffee and dark chocolate were linked to SCFA production due to their bioactive compounds, like caffeine, chlorogenic acid, and polyphenols, which modulate the gut microbiota and fermentation activities ^77, 78^. These findings emphasize the critical role of diet in regulating SCFA levels and support our model’s effectiveness in identifying dietary determinants of SCFA production.

### Host characteristics

Our study identified several factors in host characteristics—such as gender, race, age, height, weight, BMI, physical activity, and the use of probiotics, antibiotics, and gastric acid-lowering medications—were correlated with blood SCFA levels. Race was associated with SCFA production, aligning with previous findings that African Americans have lower fecal acetate levels compared to white participants ^45^, potentially reflecting similar trends in blood SCFAs ^79^. Gender and age also affect SCFA production due to differences in gut microbiota diversity and composition across groups ^80, 81^. Body composition indicators, such as BMI, weight, and height, correlate with SCFA levels, as reduced gut microbiota diversity in overweight or obese individuals often results in increased SCFA production ^82–84^, which is linked to energy storage and lipid metabolism ^85–87^. Physical activities such as biking and swimming affect muscle lactate metabolism ^88–90^, and *Veillonella* species in the gut can convert lactate into SCFAs like acetic acid ^91^. Additionally, probiotics increase SCFA production by boosting SCFA-producing bacteria ^92^, whereas antibiotics and gastric acid-lowering medications reduce microbial diversity and alter gut environments, impacting SCFA levels ^84, 93, 94^. These identified host characteristics, in turn, demonstrate the effectiveness of our model in imputing SCFA levels by integrating gut microbiome compositions, dietary features, and host characteristics, providing a comprehensive understanding of the determinants influencing SCFA production.

Our current results indicated that the regulation of blood SCFAs could be a complex procedure, the GMs can be important factors. Besides, dietary habits and host characteristics might also influence blood SCFAs directly or through interactions with GMs. However, there are a few limitations in this study. First, all the subjects in our study were Caucasians and African Americans, making it necessary yet to generalize the results to other racial populations. The validation of our model in diverse populations would enhance its applicability. This necessitates further model validation in different cohorts that incorporate metagenomic and blood SCFAs profiling, dietary features, and host characteristics of a similar scope. Second, our study utilized two different datasets for model validation: one with serum SCFA measurements and the other with plasma SCFA measurements. While previous studies suggest that SCFA levels are generally consistent between serum and plasma samples, minor variations might still exist due to the different biological matrices. These variations were not directly analyzed in our study, as the primary goal was to validate the model’s performance across different datasets rather than compare serum and plasma SCFA levels. To strengthen the model’s accuracy and reliability, future studies should validate our models using additional datasets with more serum and plasma SCFA measurements. Third, another limitation of our study is the sex distribution across the two datasets used for model validation. Dataset 1 included males, while Dataset 2 included both males and females. This imbalance in sex representation between the two datasets could affect the model’s ability to generalize across different sexes. However, because Dataset 2 has a relatively small number of male participants, splitting this dataset by sex for separate analyses would result in low statistical power, leading to potentially unreliable results. To maintain robust model validation, we combined the male and female samples in Dataset 2, which helps to preserve adequate sample size for analysis. We adjusted for sex as a covariate in our model to account for potential sex-specific differences in SCFA production. However, future studies with more balanced sex distributions or larger sample sizes for both sexes would provide a more comprehensive understanding and enhance the robustness of the model. Fourth, the integration of data from two different sequencing platforms in Dataset 1 could be considered a limitation. However, deep learning models are particularly capable of finding common patterns across heterogeneous data by learning generalizable features that are not specific to any single sequencing platform. We applied uniform normalization within the model to standardize the data and employed regularization techniques like dropout and mean absolute error loss to prevent overfitting to sequencing platform-specific characteristics. While these strategies allow the model to generalize effectively, future studies could include more data generated from either consistent or varied sequencing platforms to further validate the model’s robustness across different technical conditions. Fifth, another limitation of our study is that the features in each view of the two datasets differ, which could result in variability in the important features identified by the model. Although many of the identified important features are well-known factors related to SCFA production, demonstrating the model’s effectiveness to some extent, this variability highlights the need for caution in interpreting results. However, it is worth noting that the imputation performance was strong across both datasets, highlighting the model’s generalizability despite these differences. It also opens up new opportunities to explore and identify additional relevant features that might be specific to certain datasets or conditions. Future studies should aim to include datasets with consistent feature sets across all views to enhance comparability and validate the model’s ability to generalize findings across different contexts.

To summarize, we have blazed a trail with our innovative method that synthesizes information from the gut microbiome, dietary features, and host characteristics to perform multi-view imputation for blood SCFAs data. This can also help us identify key factors or pathways that regulate blood SCFAs in the future study. Our research highlights the utility of integrating information on gut microbiome, dietary features, and host characteristics, providing fresh perspectives on the potential regulatory mechanisms affecting blood SCFAs.

## Data Availability

The raw data presented in this study can be found in online repositories. The names of the repositories and accession numbers can be found below: NCBI BioProjects PRJNA1015234 and PRJNA1015228.

## Code Availability

The source code of this work can be downloaded from GitHub (https://github.com/Wonderangela123/M2AE).

## Acknowledgement

This work is made possible with partial support by grants from the NIH (U19AG055373, R01AG061917, R01AG068232, P20GM109036 and P20GM103629).

---

**Supplementary Table 1.**
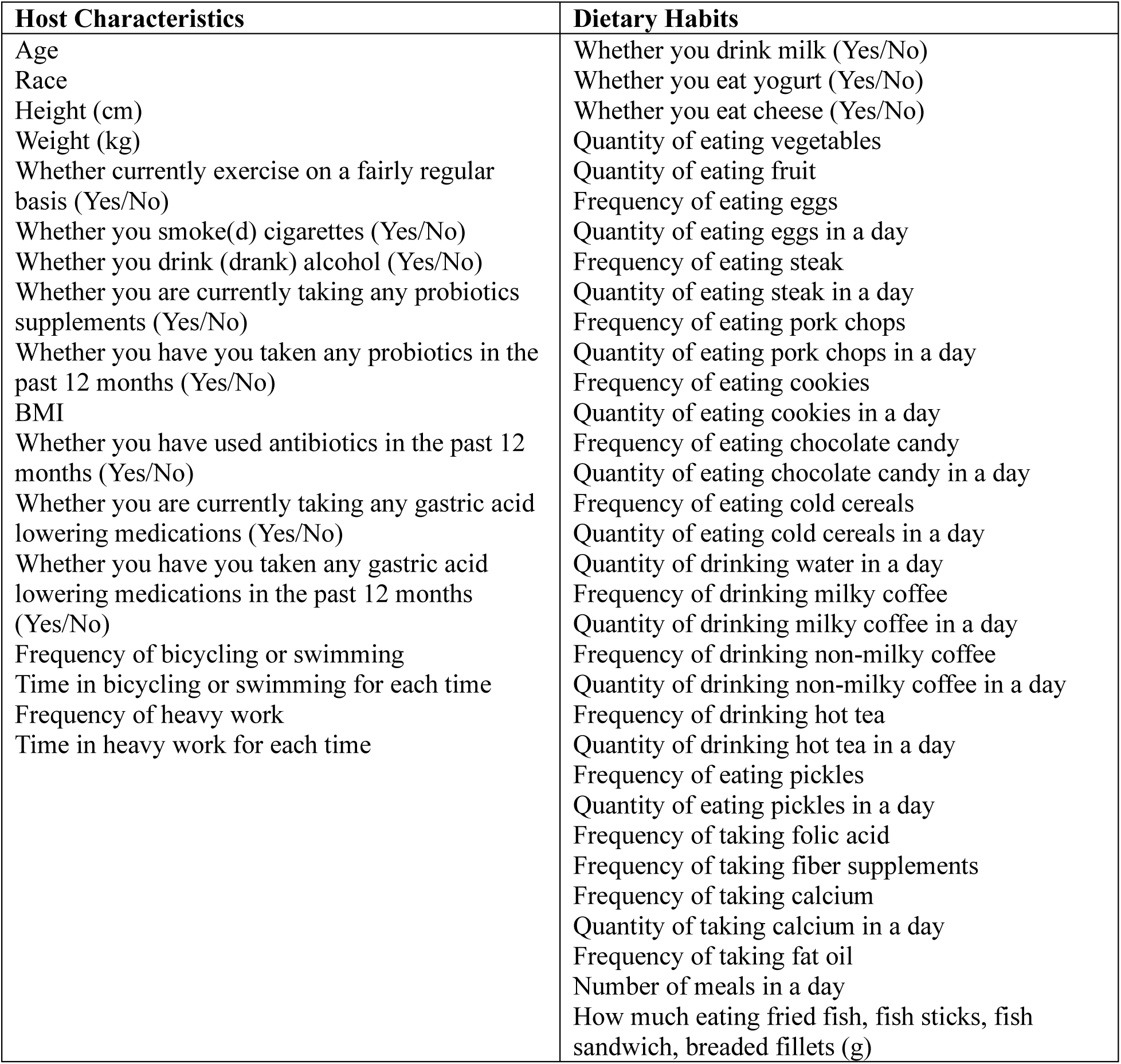
Features (Host characteristics and dietary habits) used in Dataset 1.

**Supplementary Table 2.**
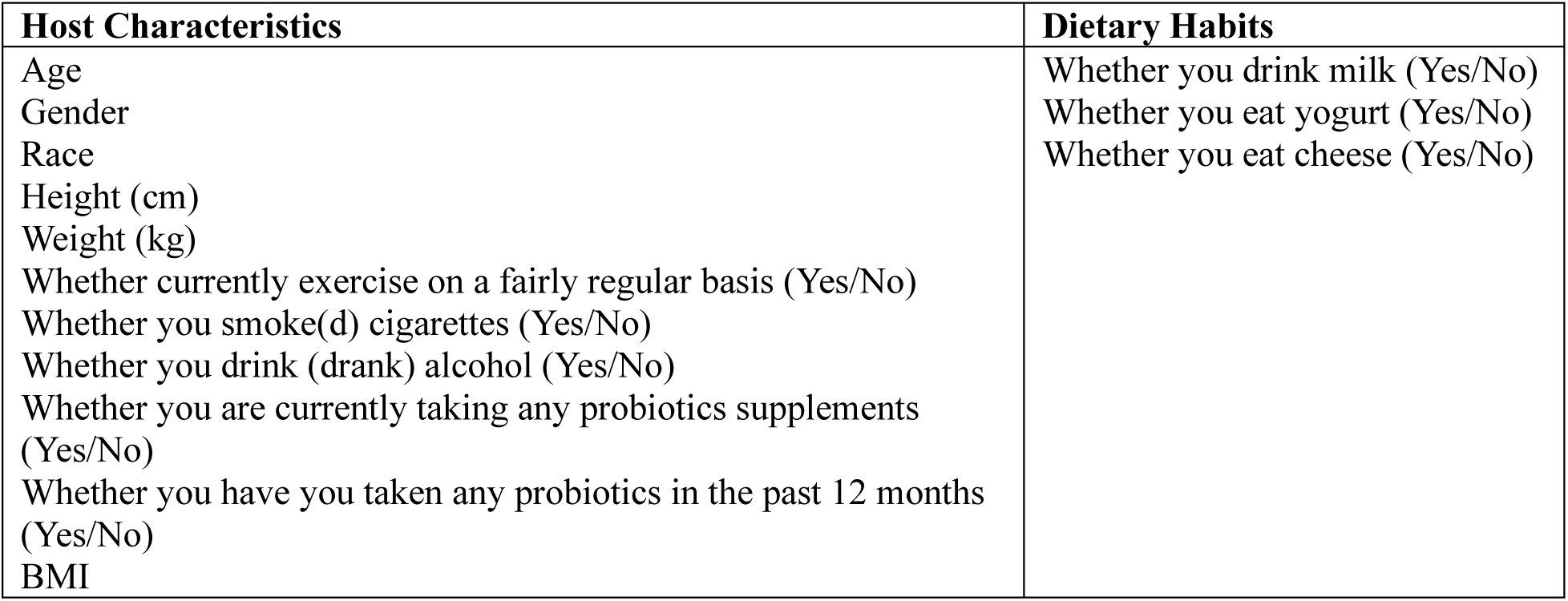
Features (Host characteristics and dietary habits) used in Dataset 2.

**Supplementary Table 3.** Hyperparameters Tuning.

| Hyperparameters | Values |
| --- | --- |
| Dropout | 0.1, 0.3, 0.5, 0.7 |
| Number of epochs | 1000, 1500, 2000, 2500, 3000 |
| Average number of edges per<br>node that are retained<br>including self-connections (k) | 2, 5, 10 |

**Supplementary Table 4.**
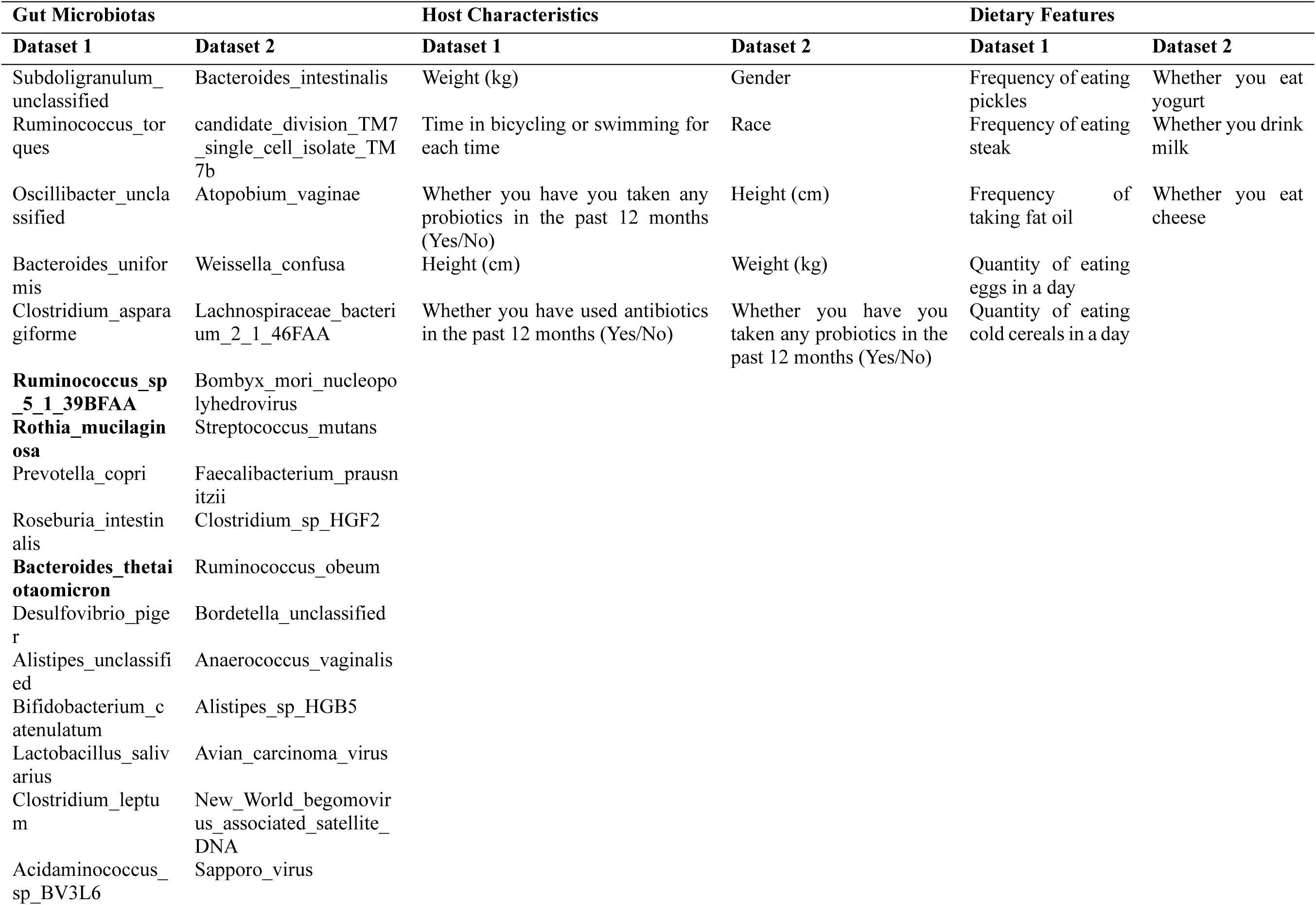
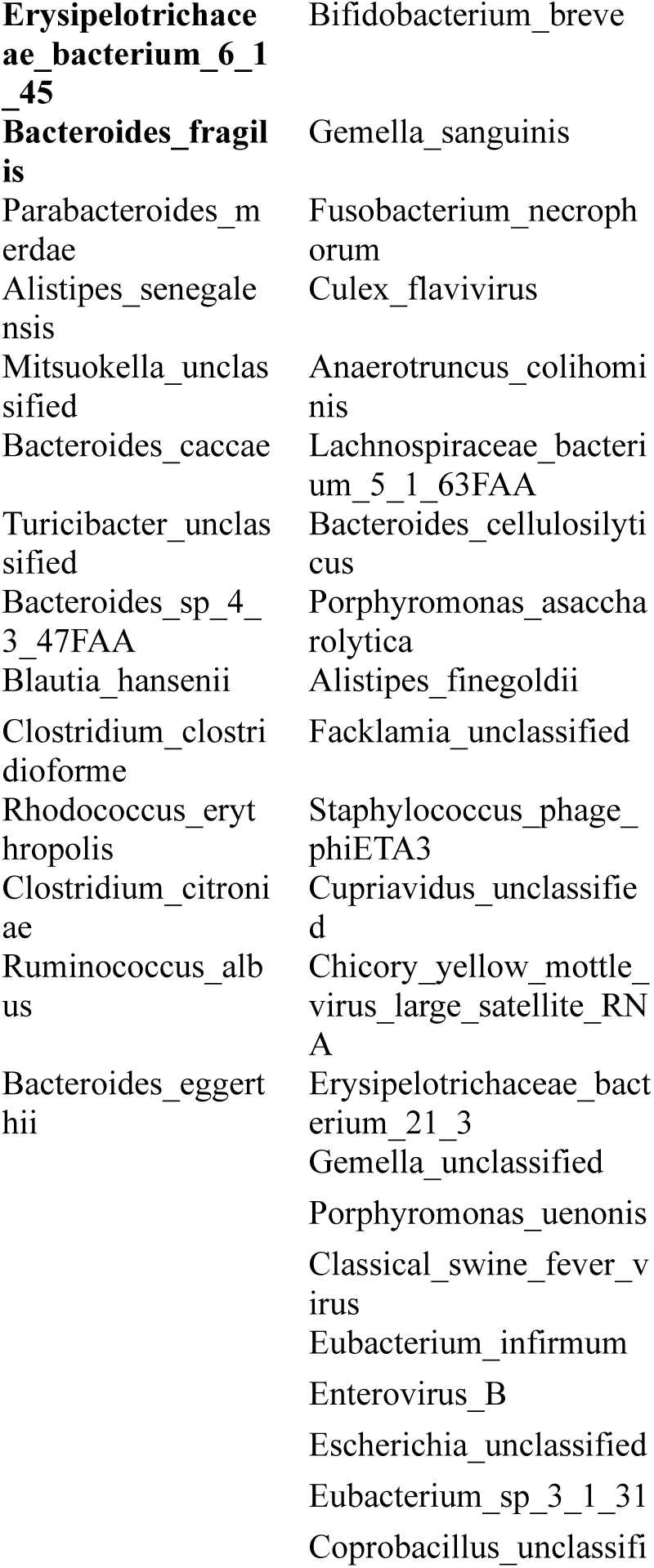
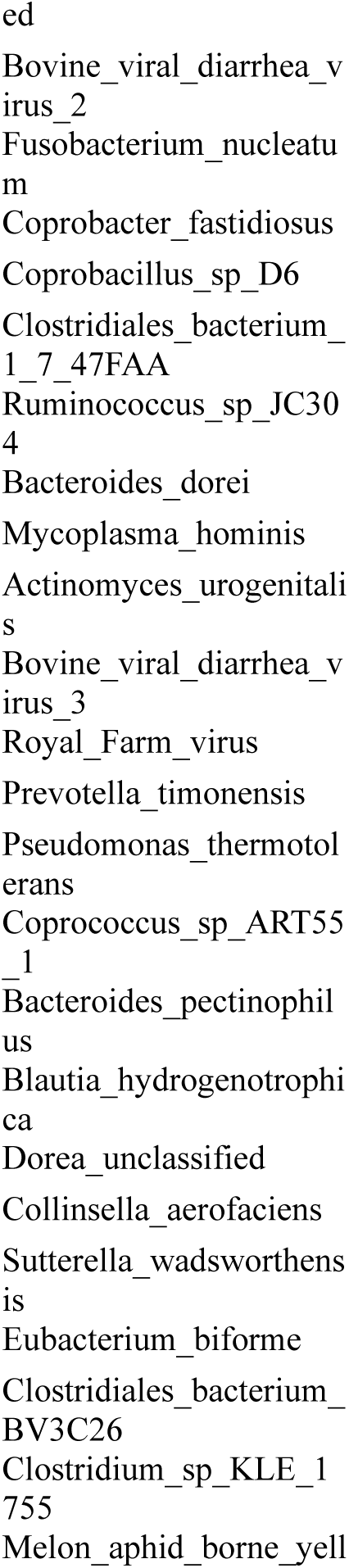
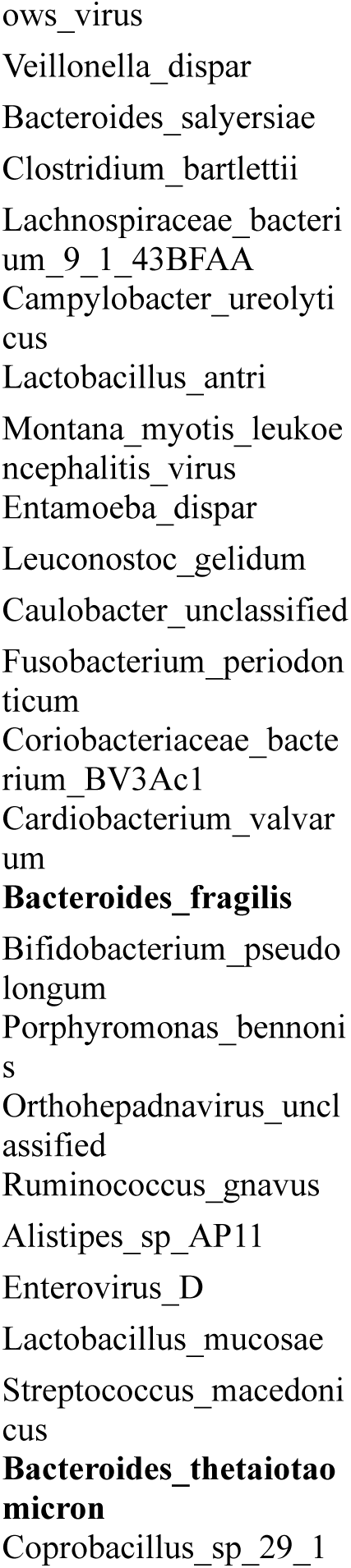
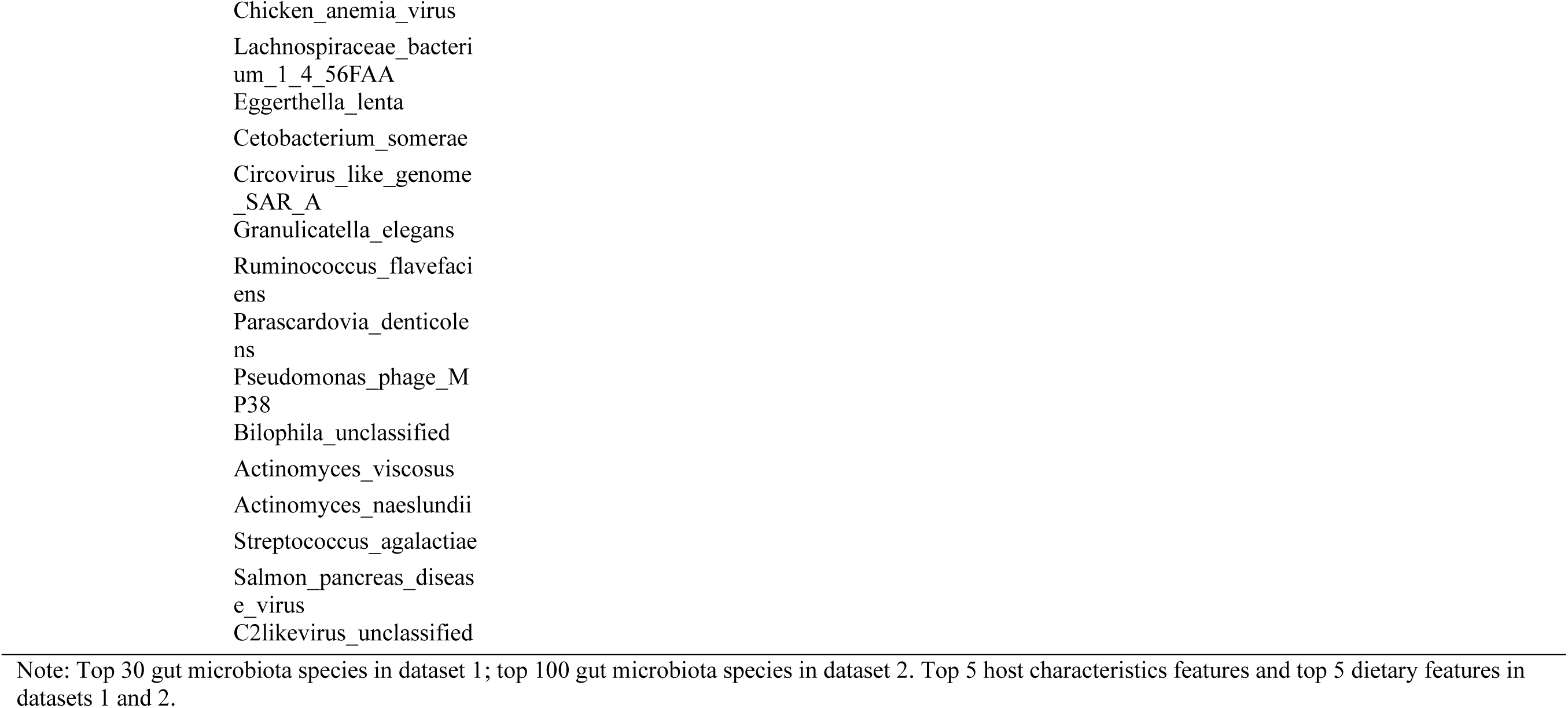
Important factors associated with blood acetic acids.

**Supplementary Table 5.**
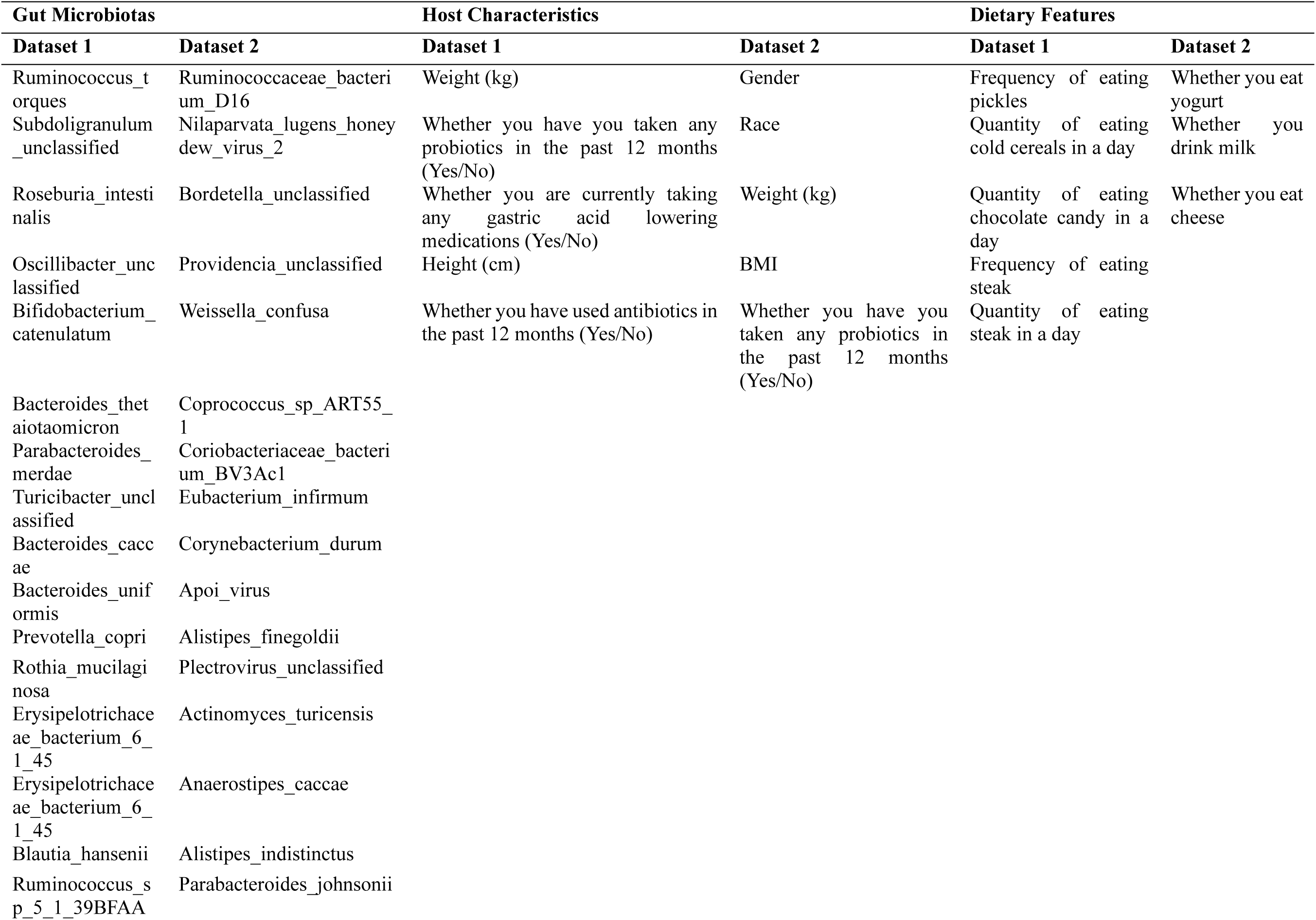
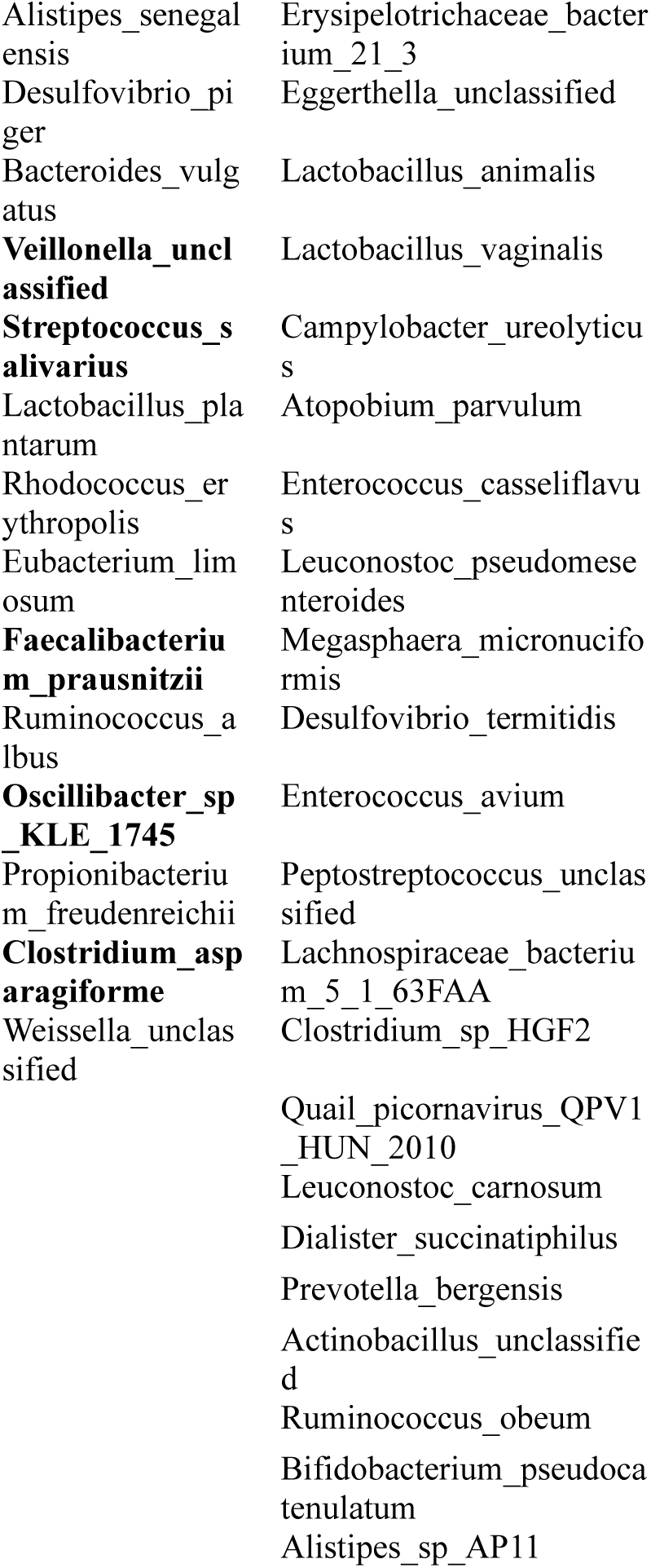
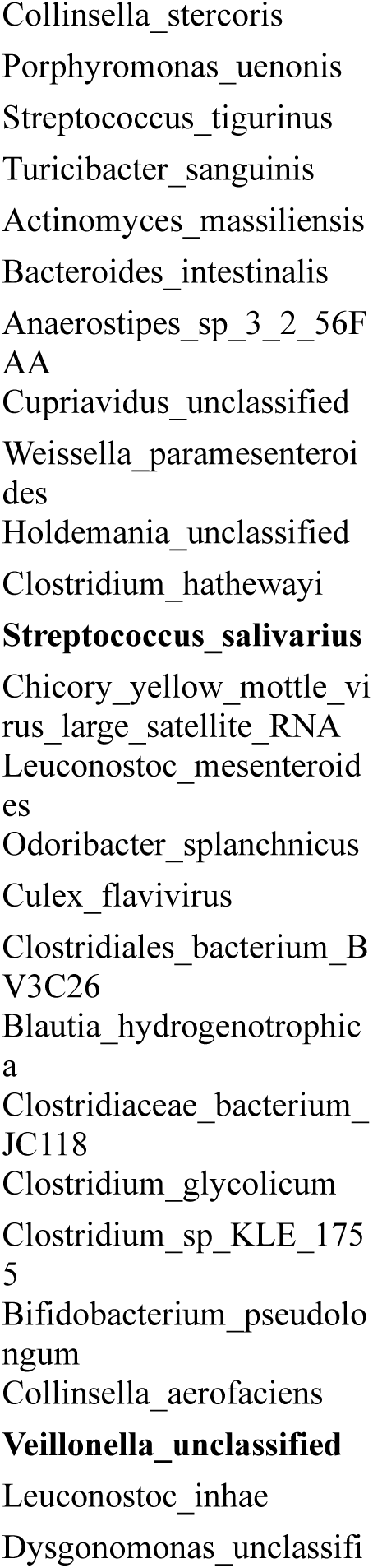
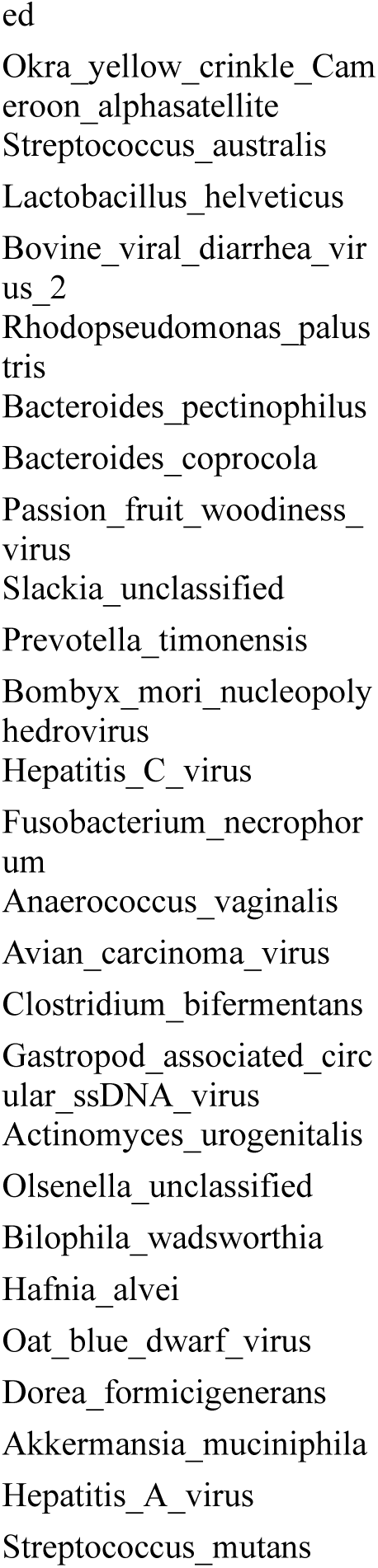
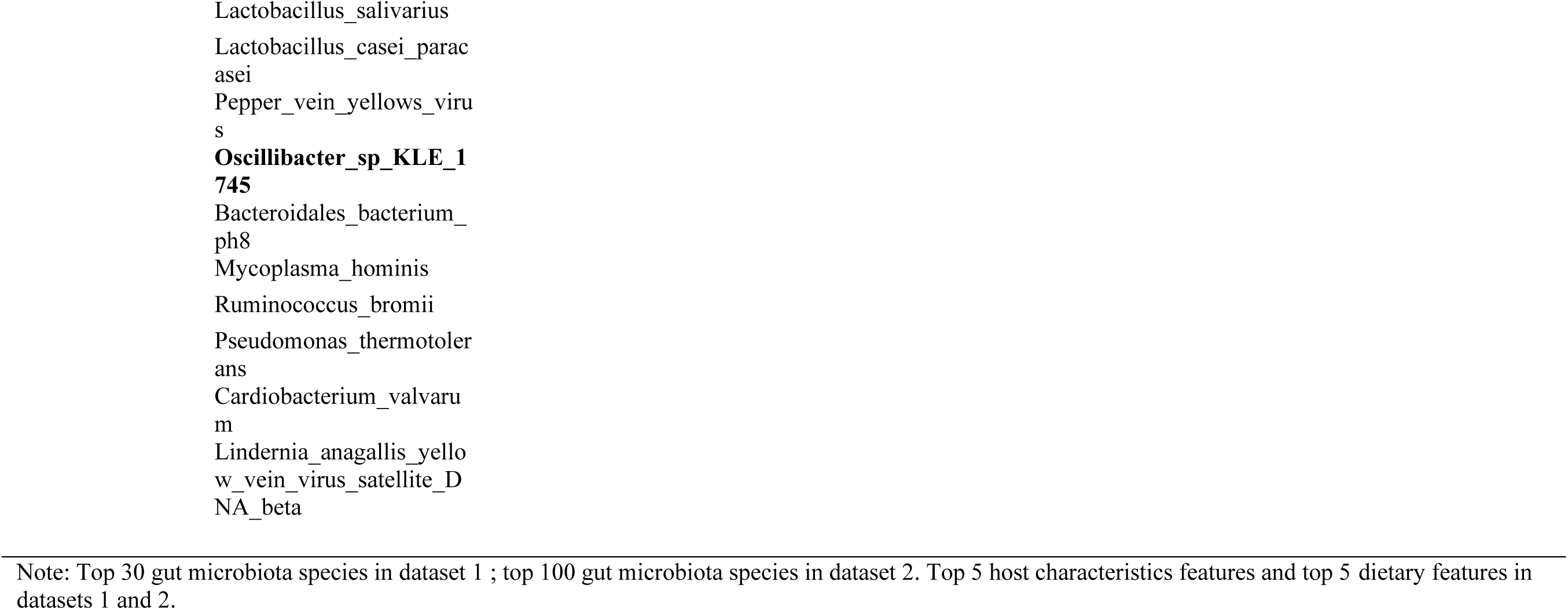
Important factors associated with blood butyric acids.

**Supplementary Table 6.**
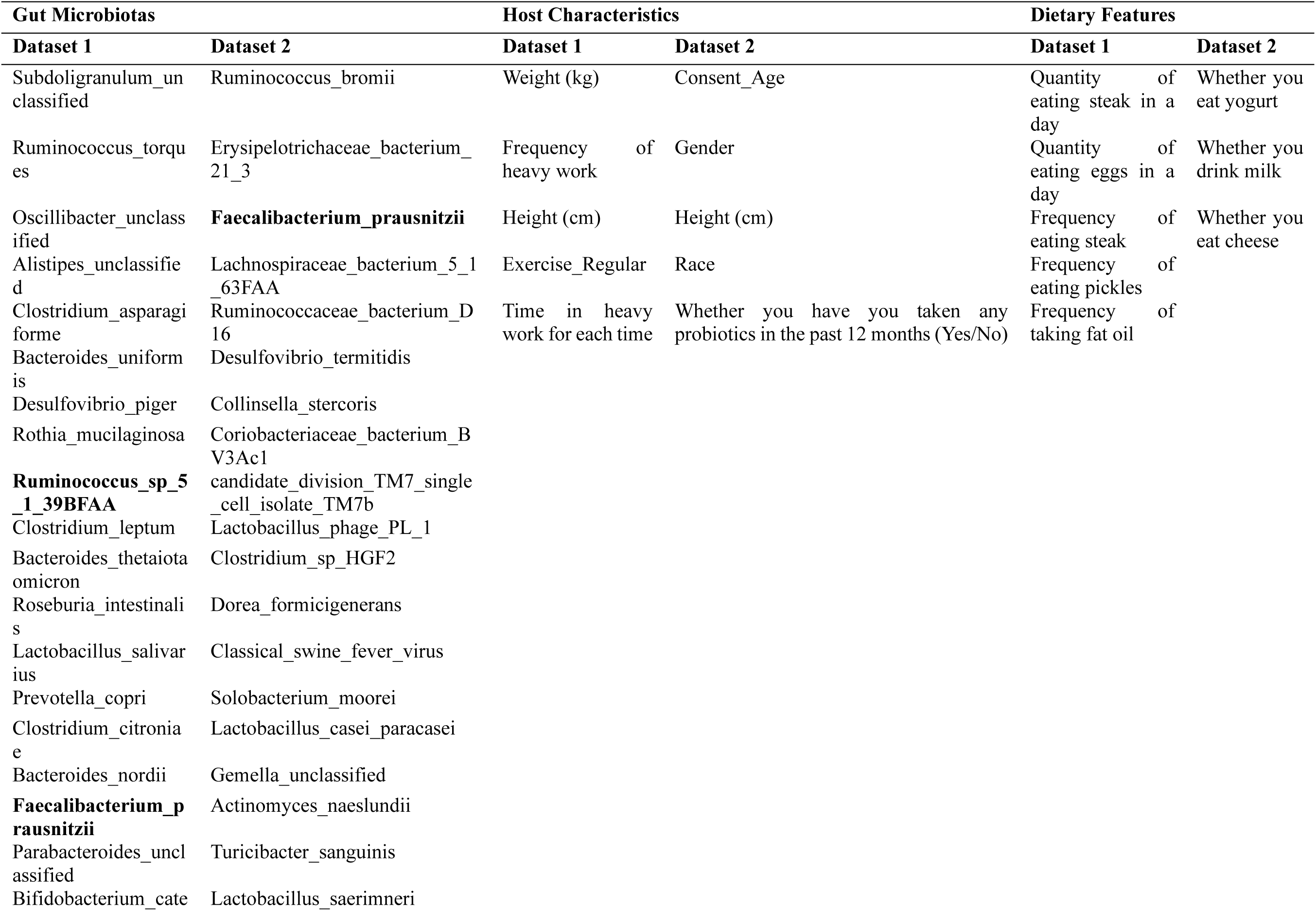
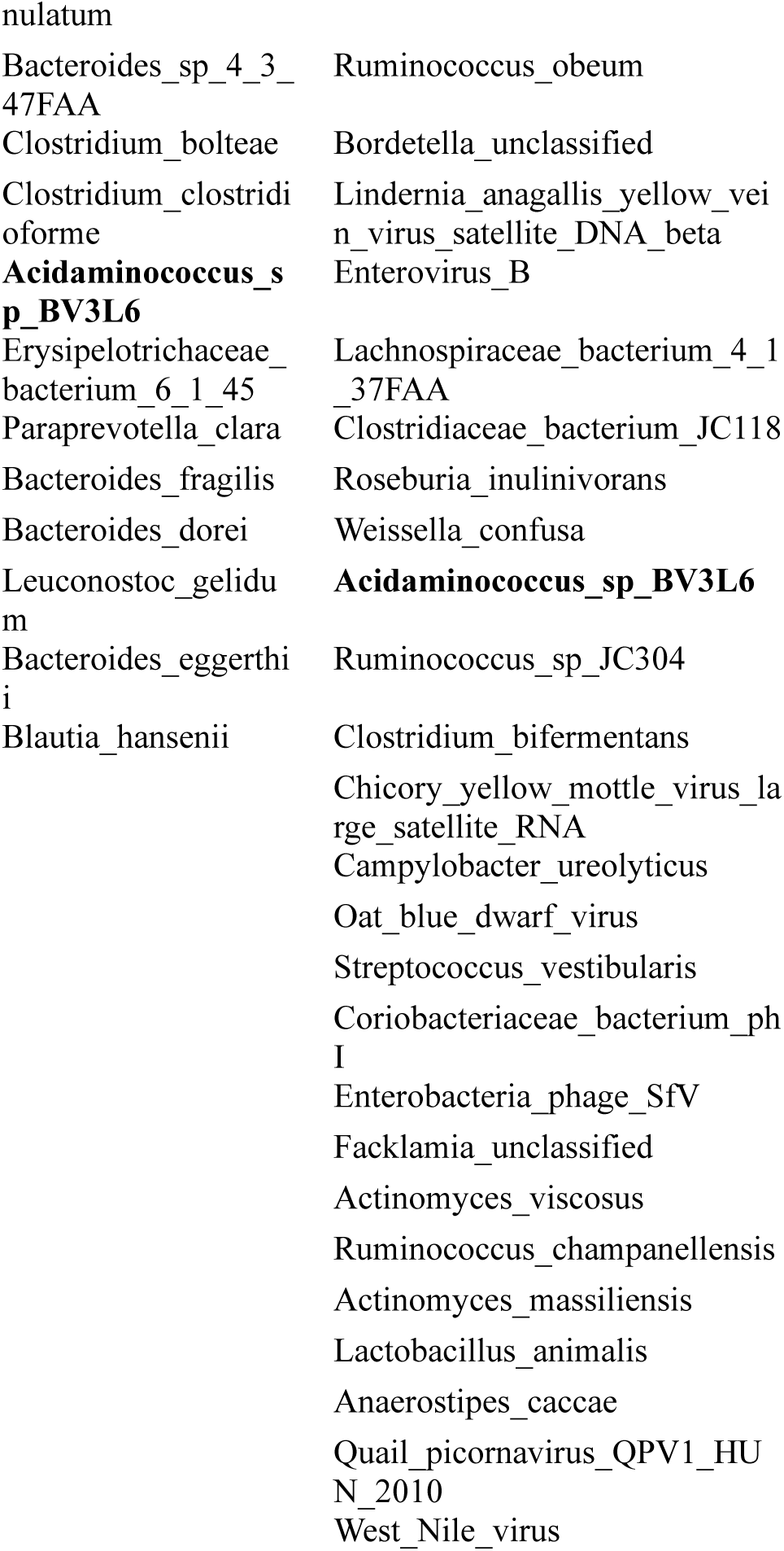
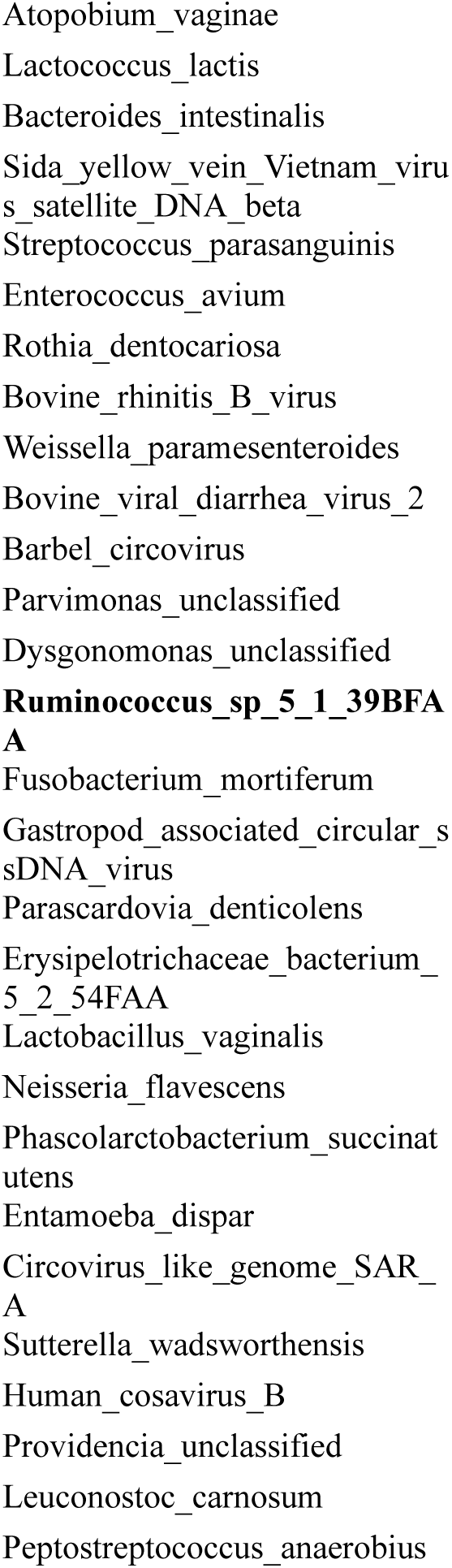
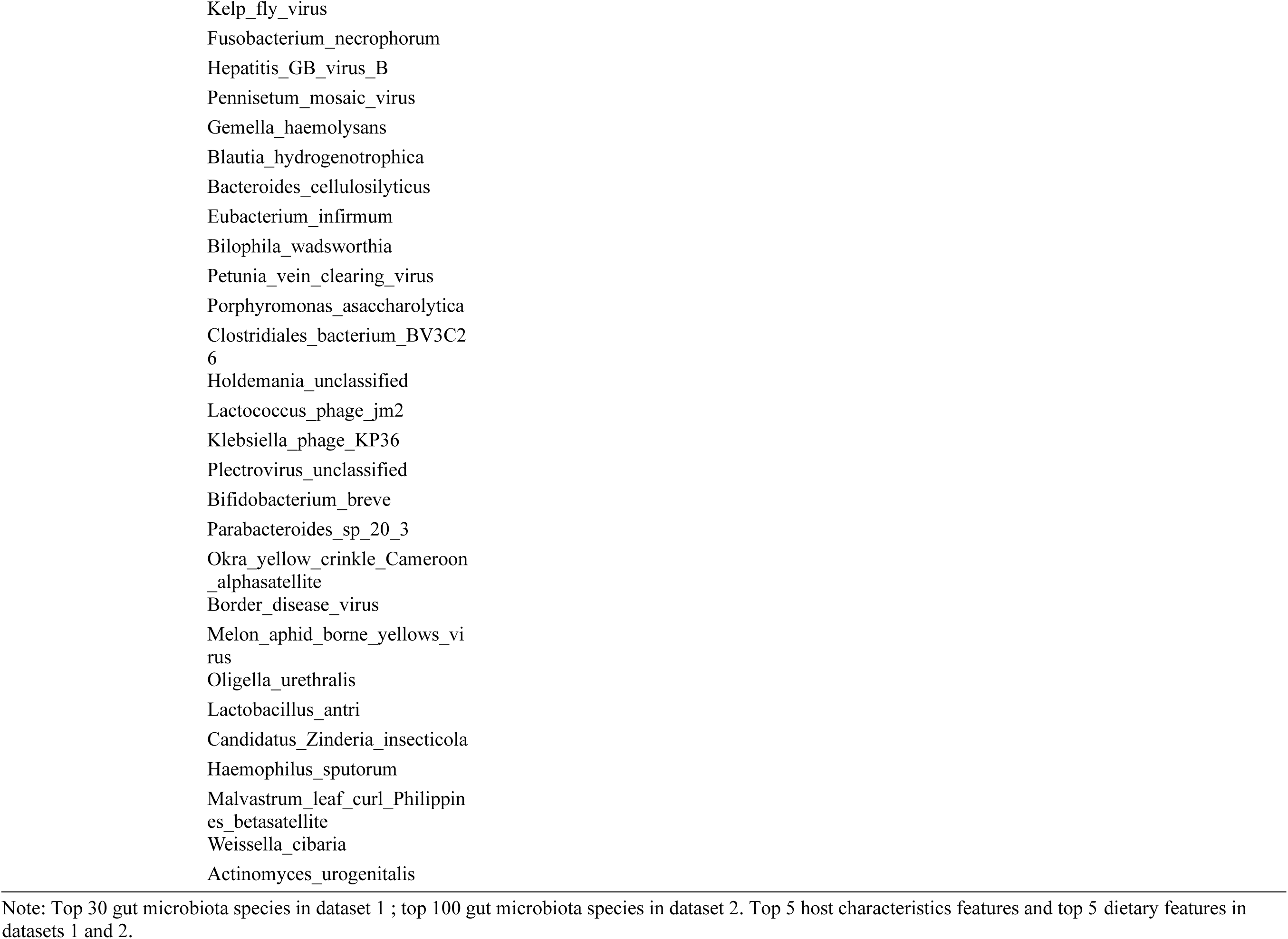
Important factors associated with blood hexanoic acids.

**Supplementary Table 7.**
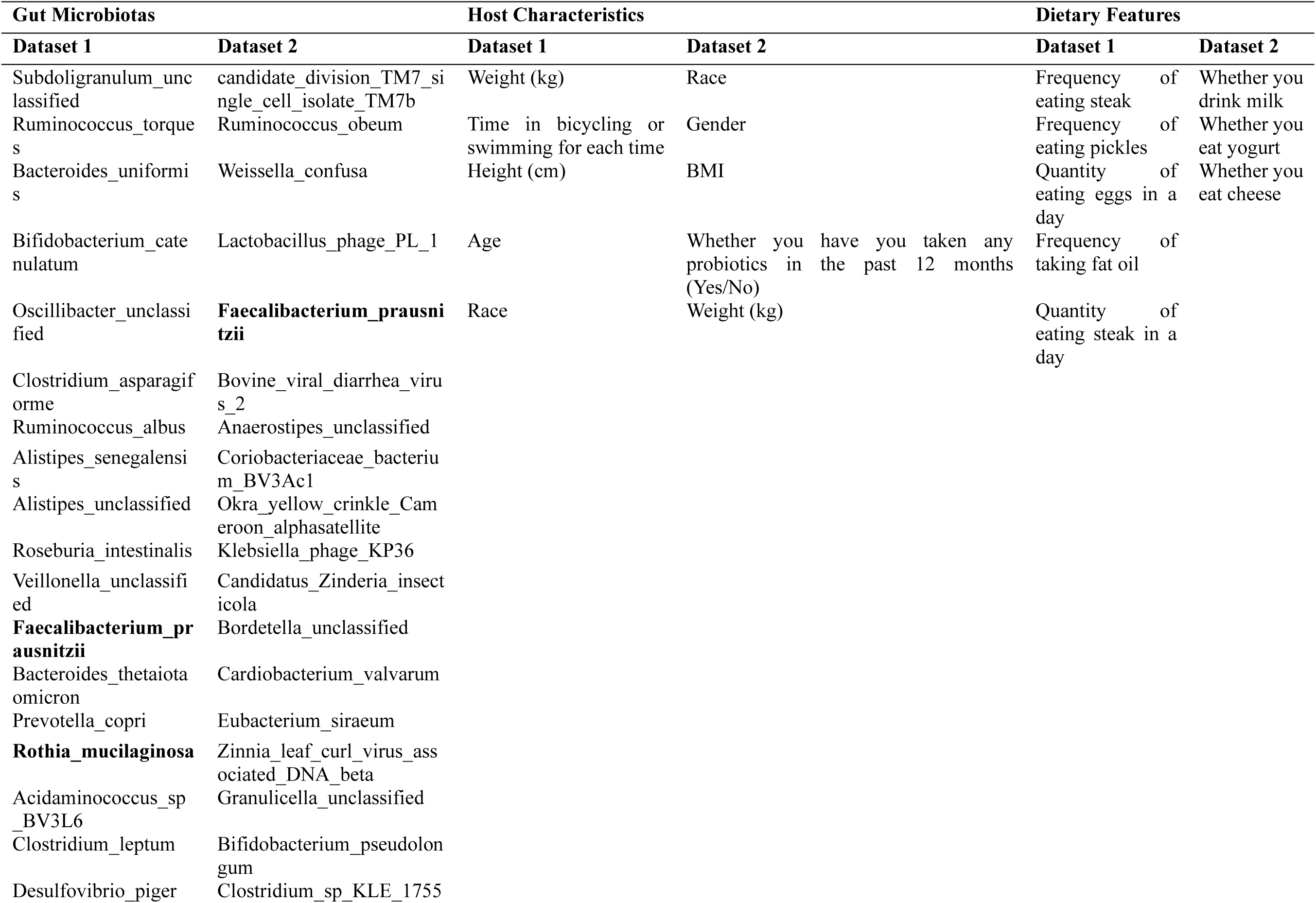
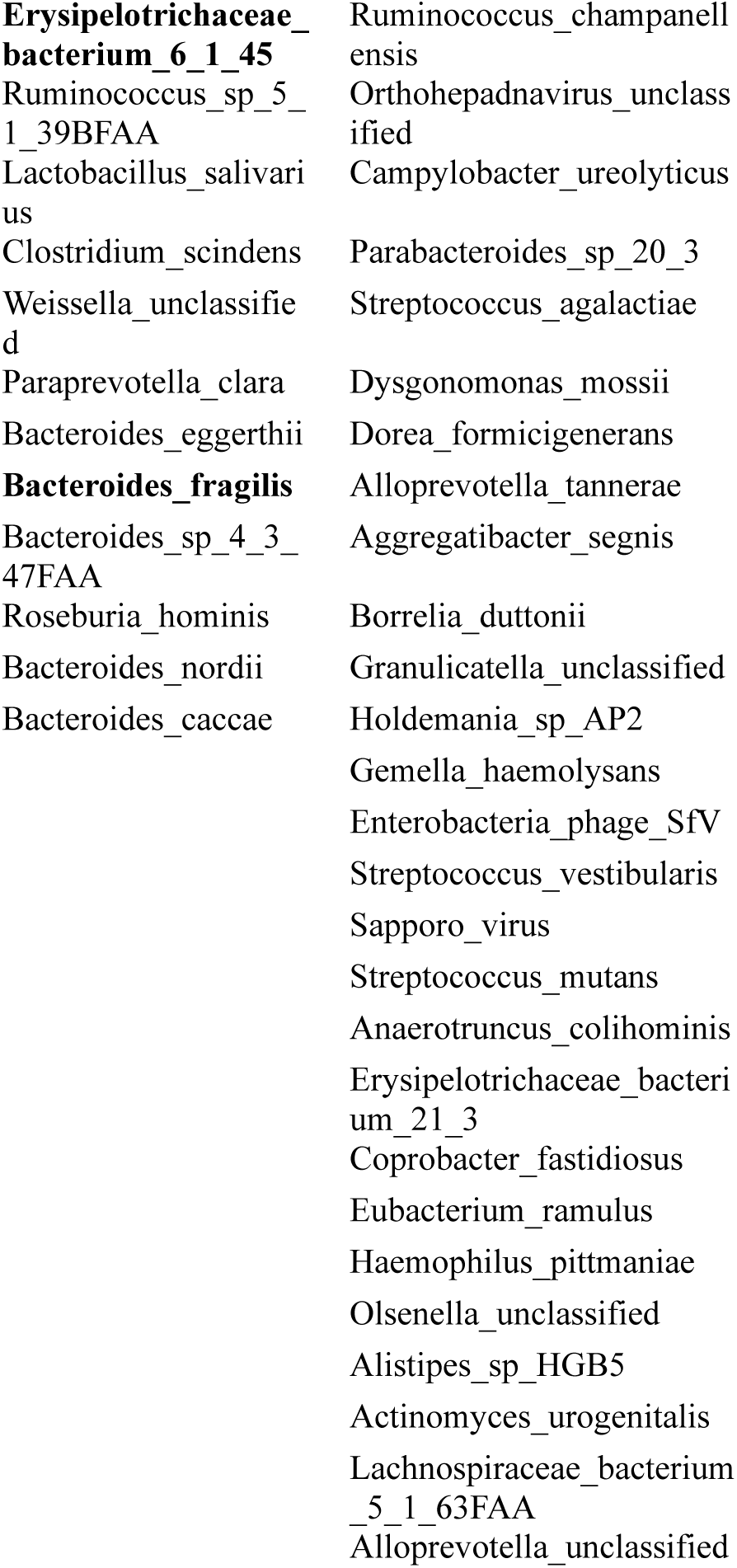
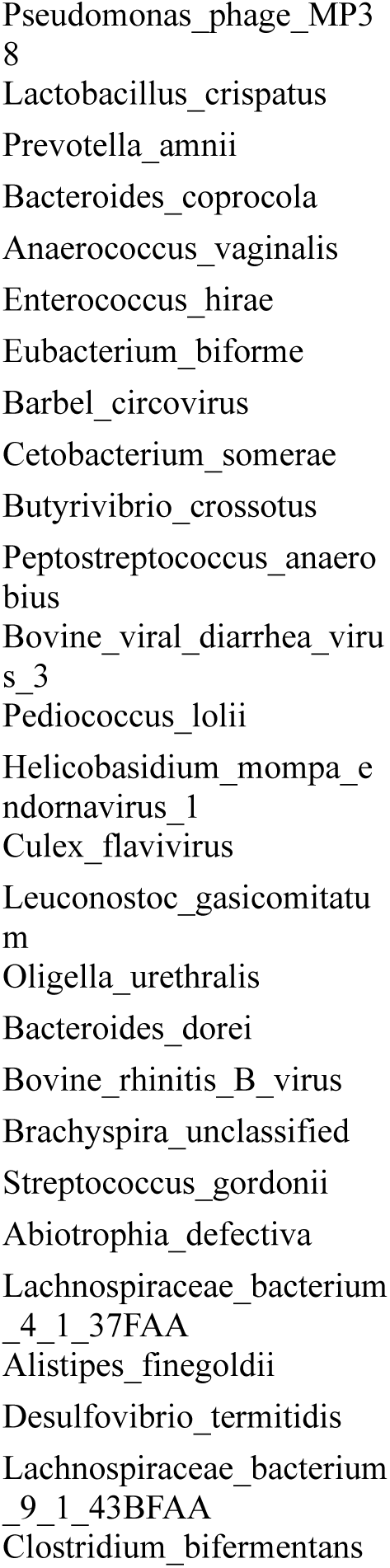
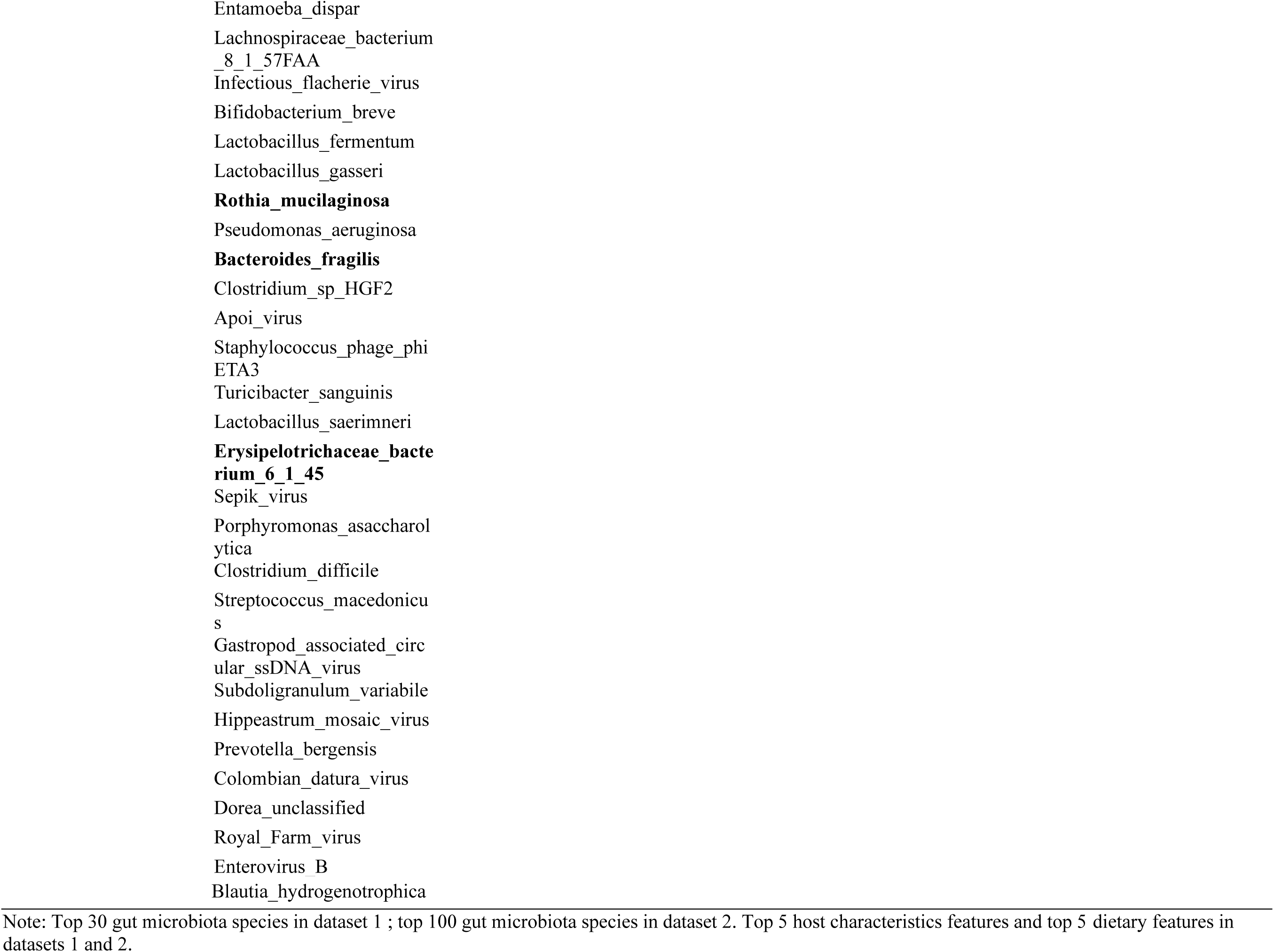
Important factors associated with blood 2-methylbutyric acids.

**Supplementary Table 8.**
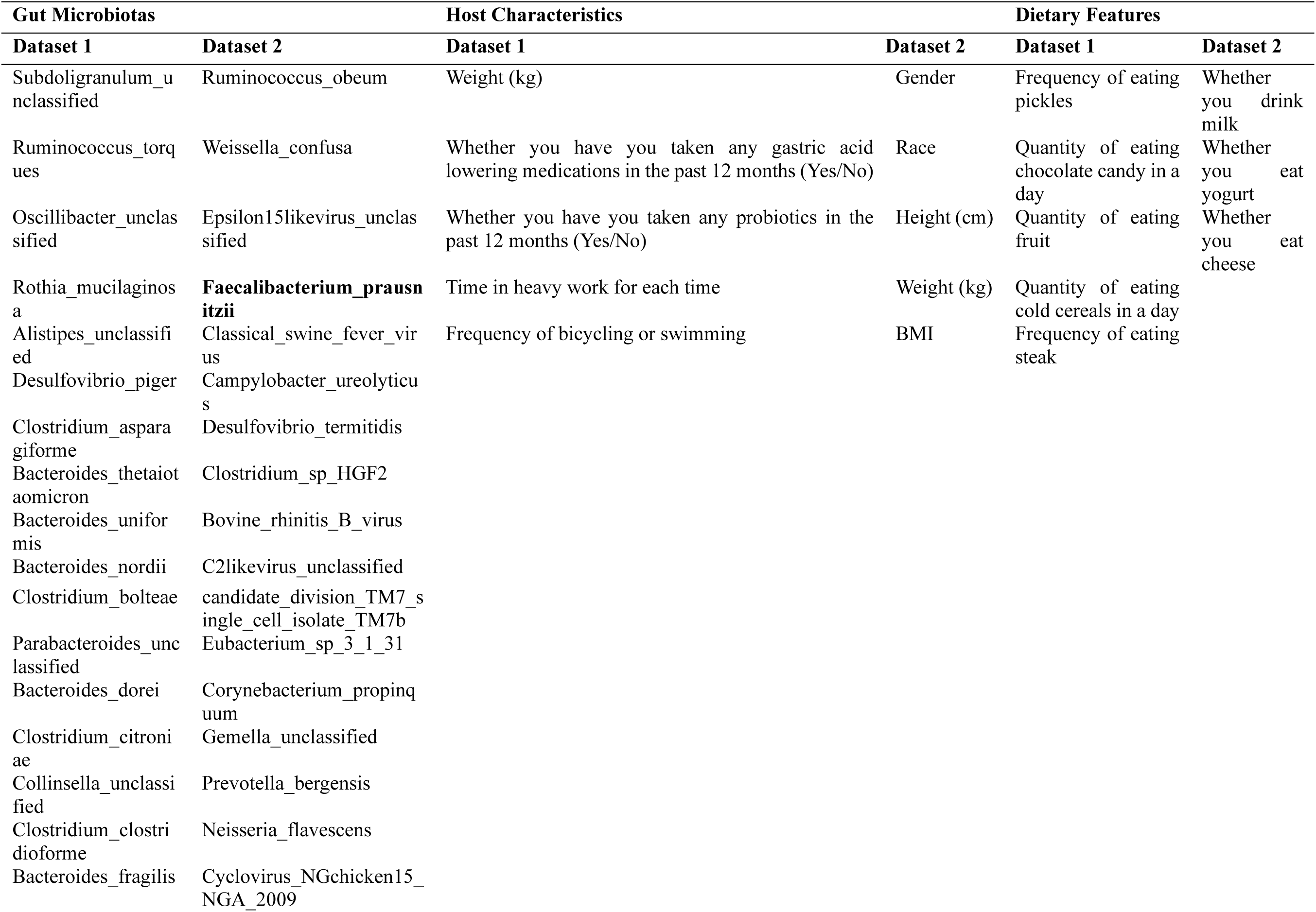
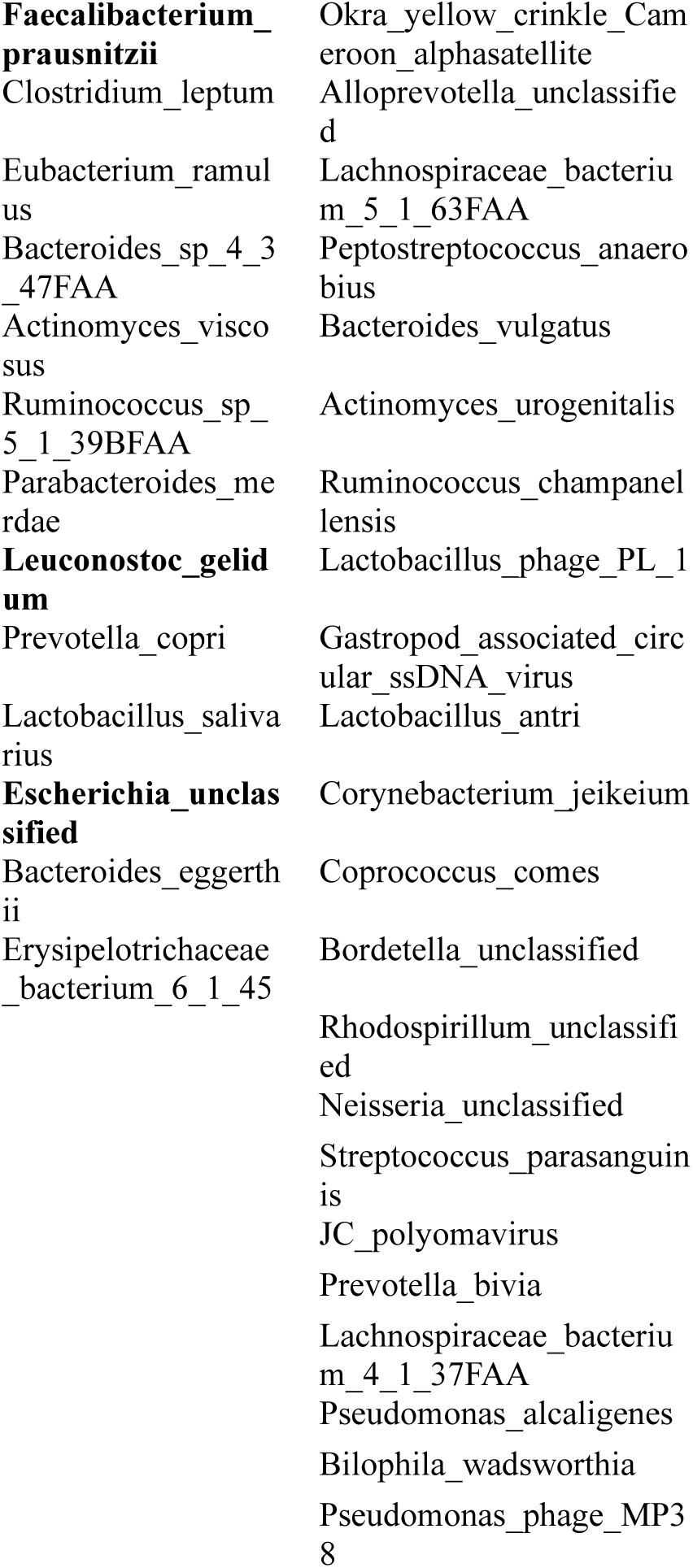
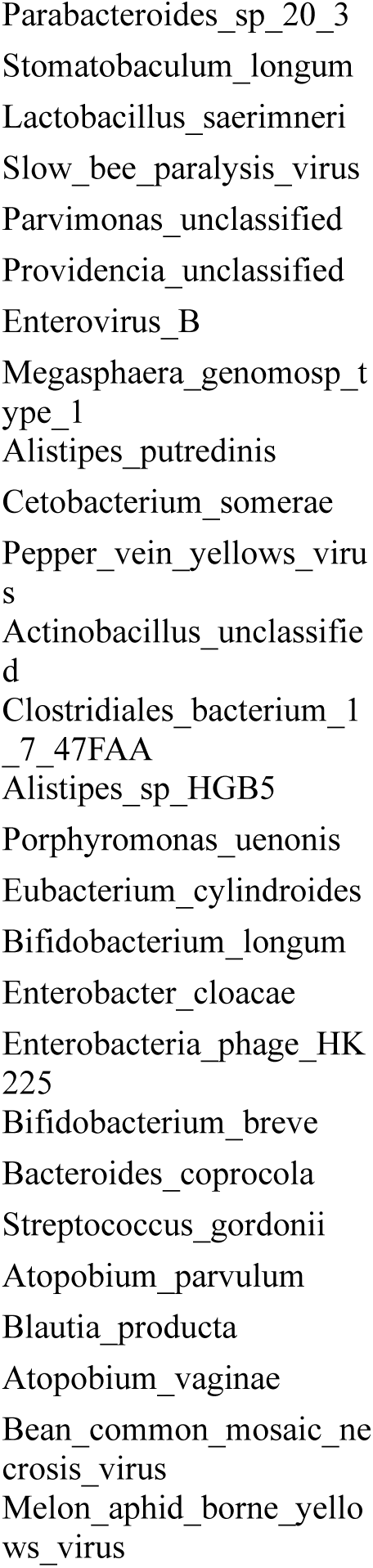
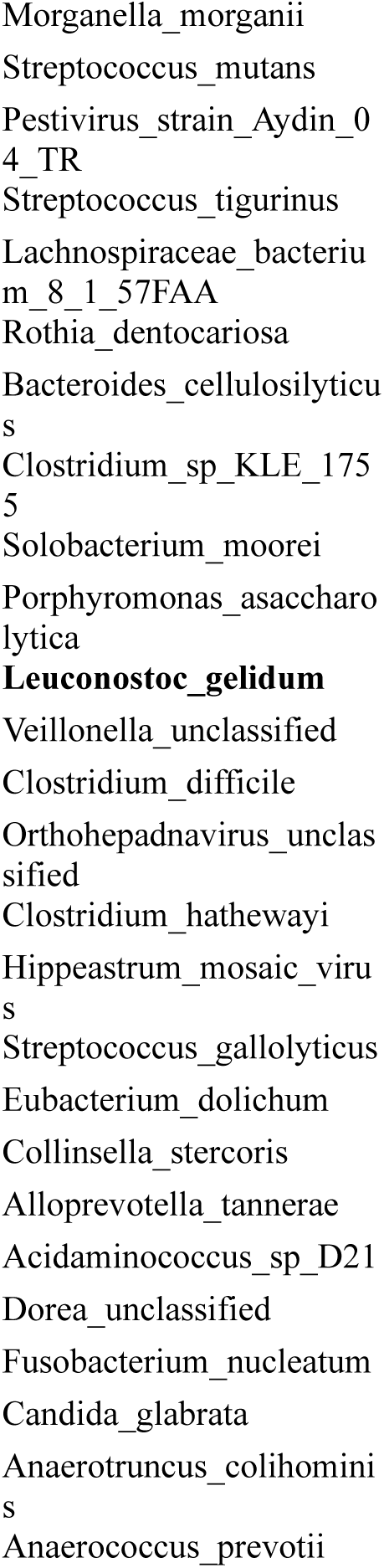
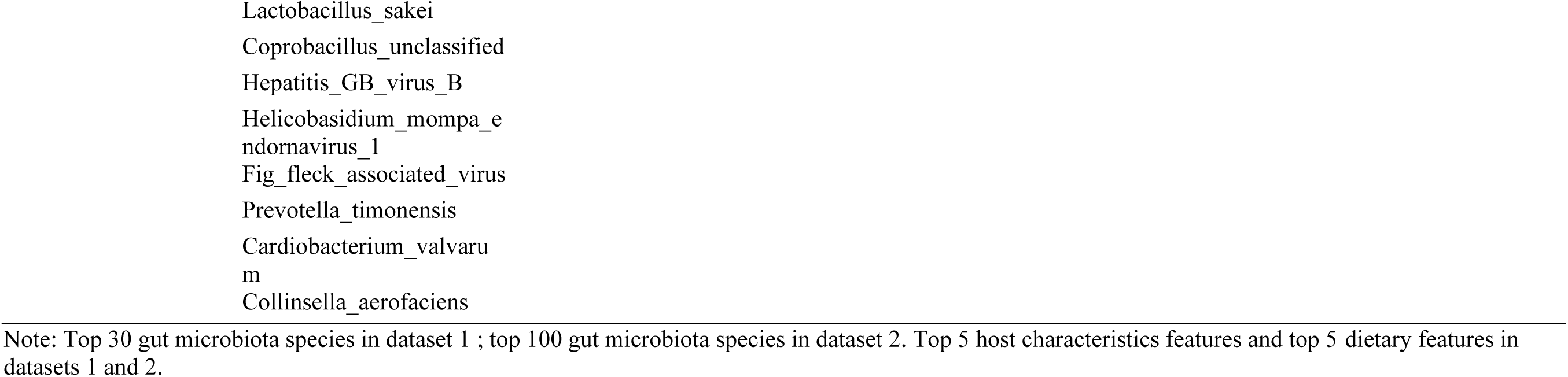
Important factors associated with blood valeric acids.

**Supplementary Table 9.**
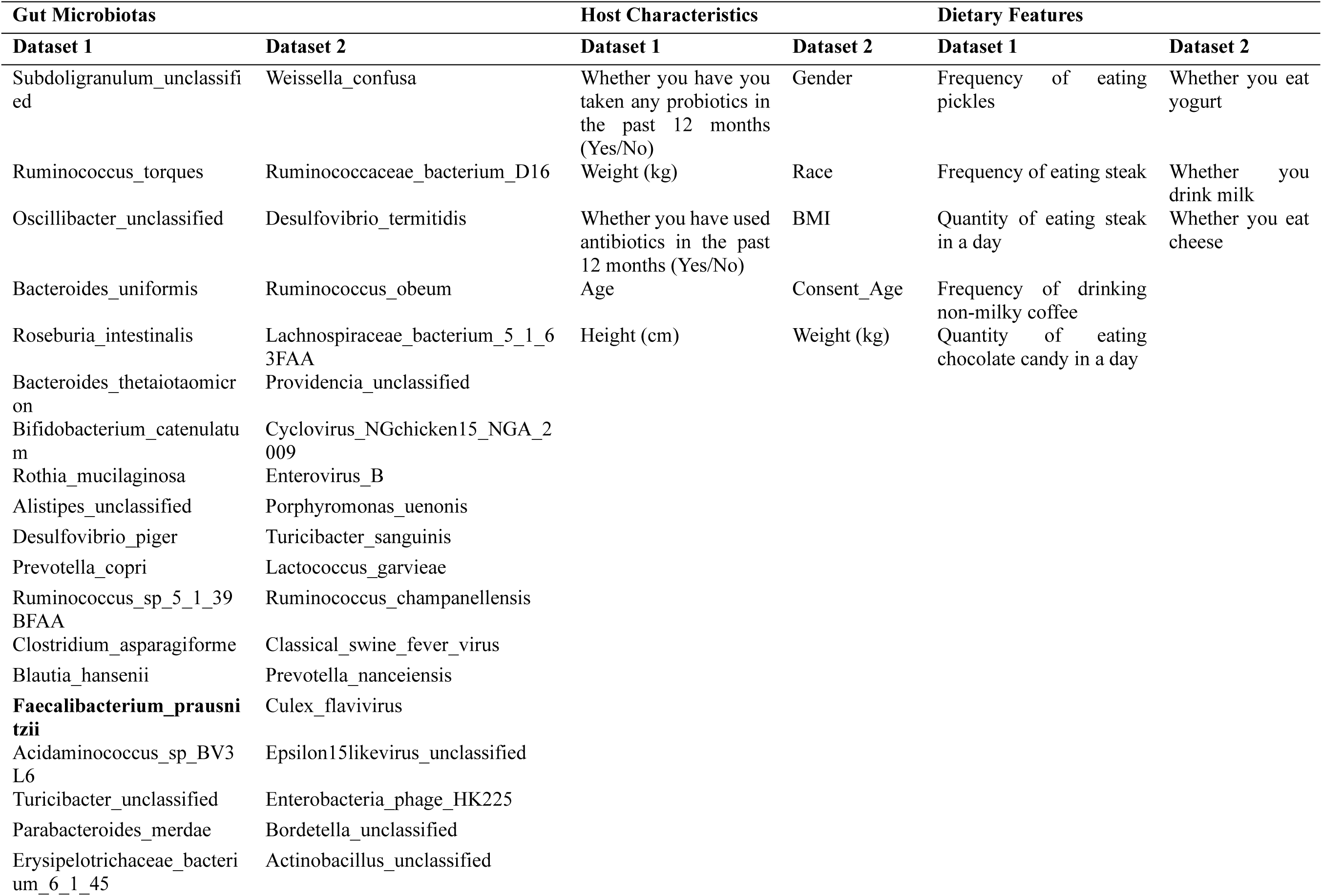
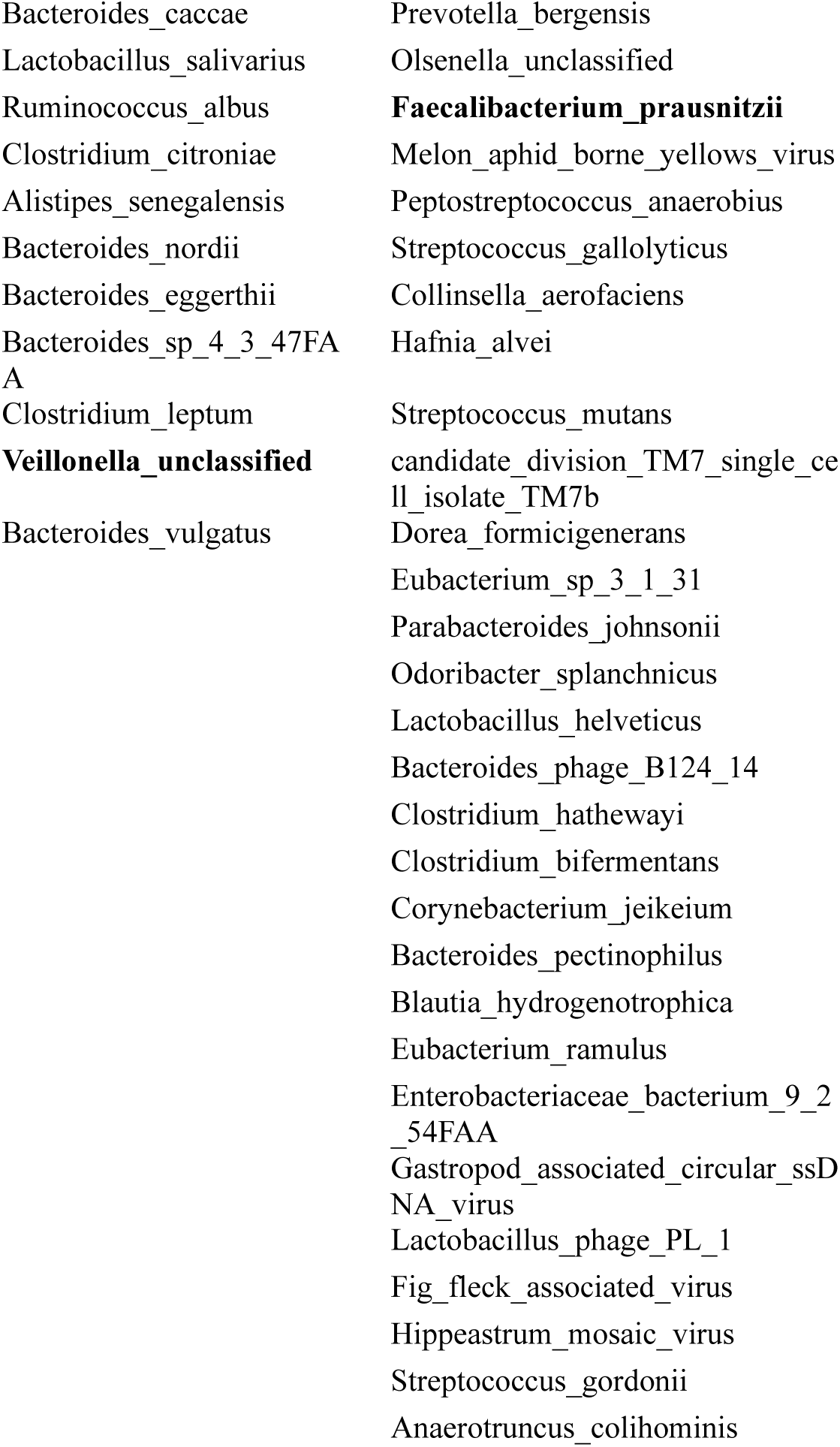
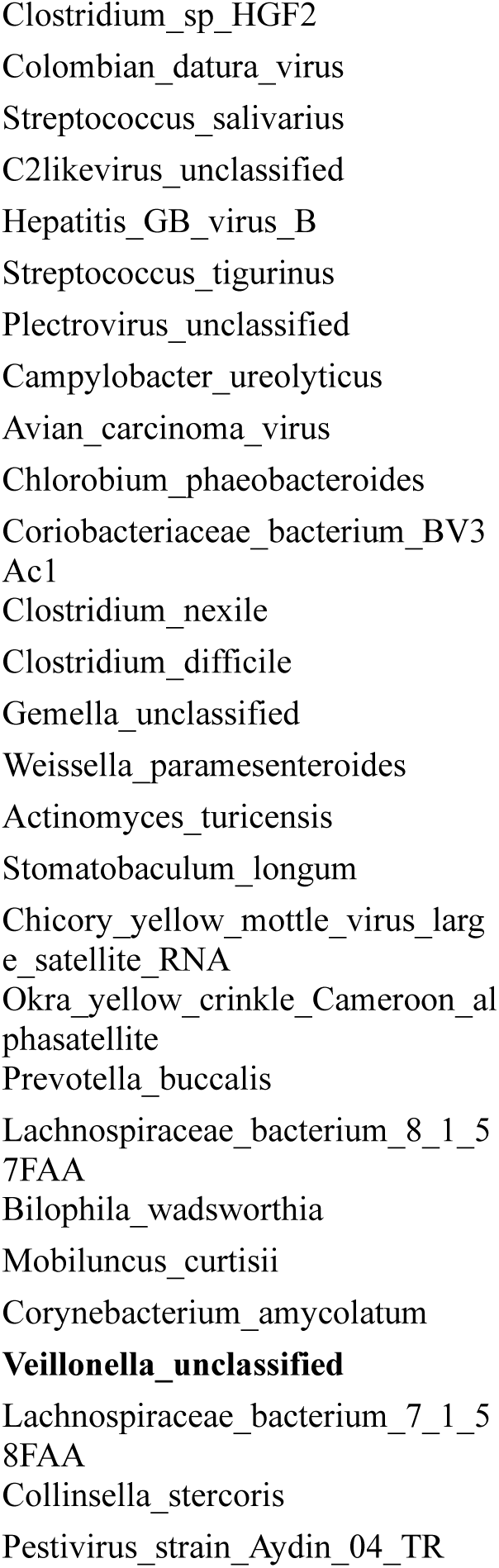
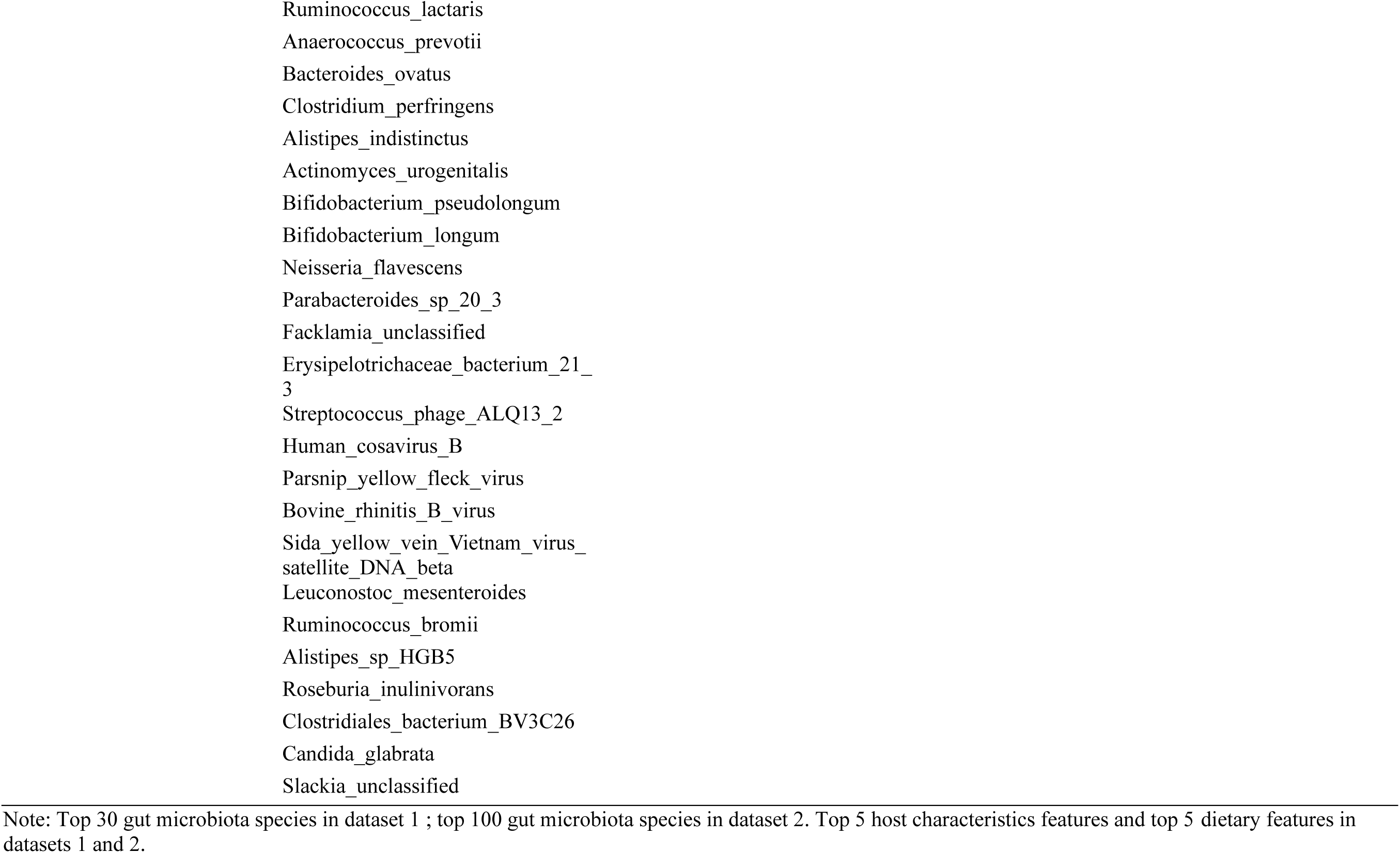
Important factors associated with blood propionic acids.

**Supplementary Table 10.**
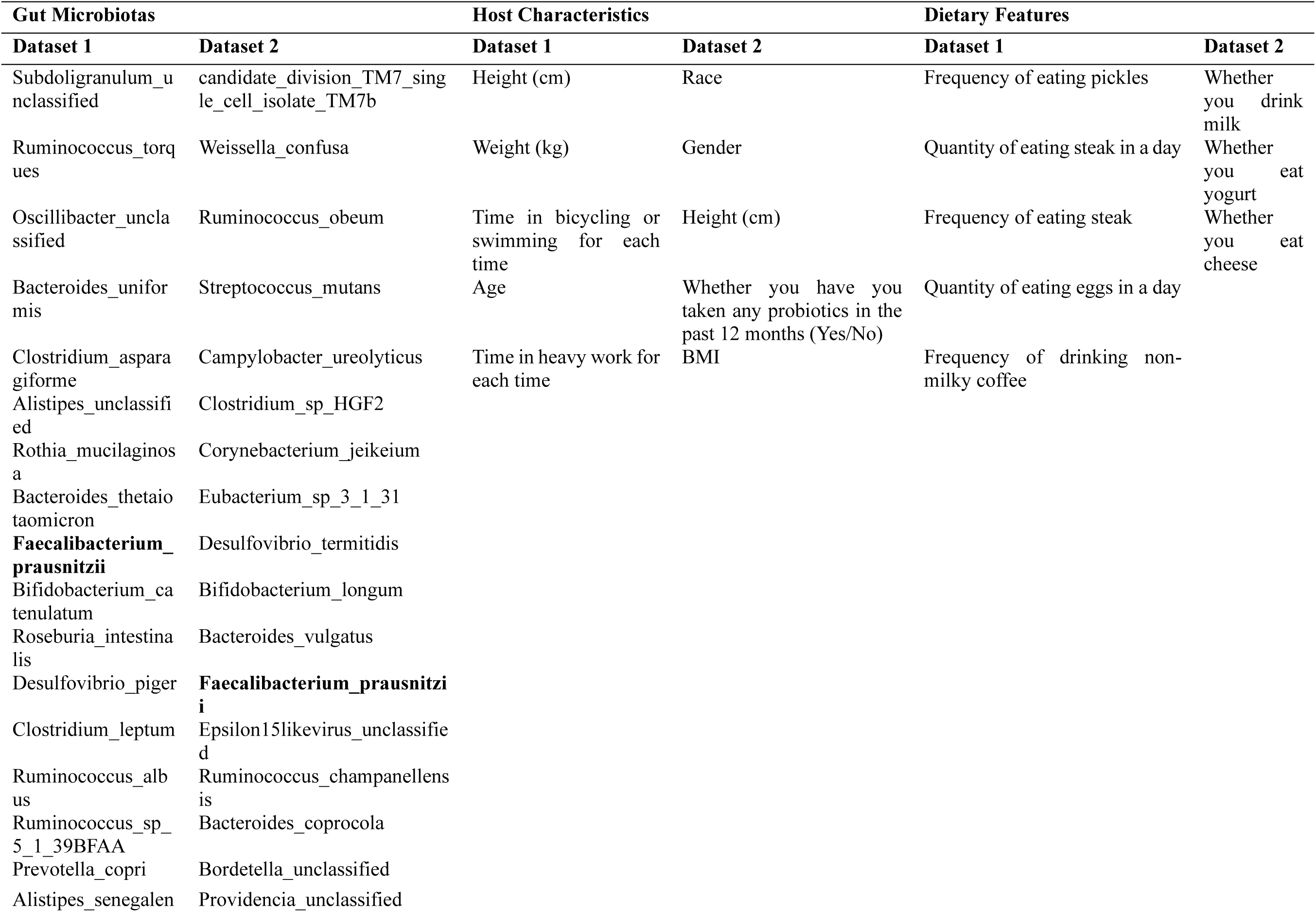
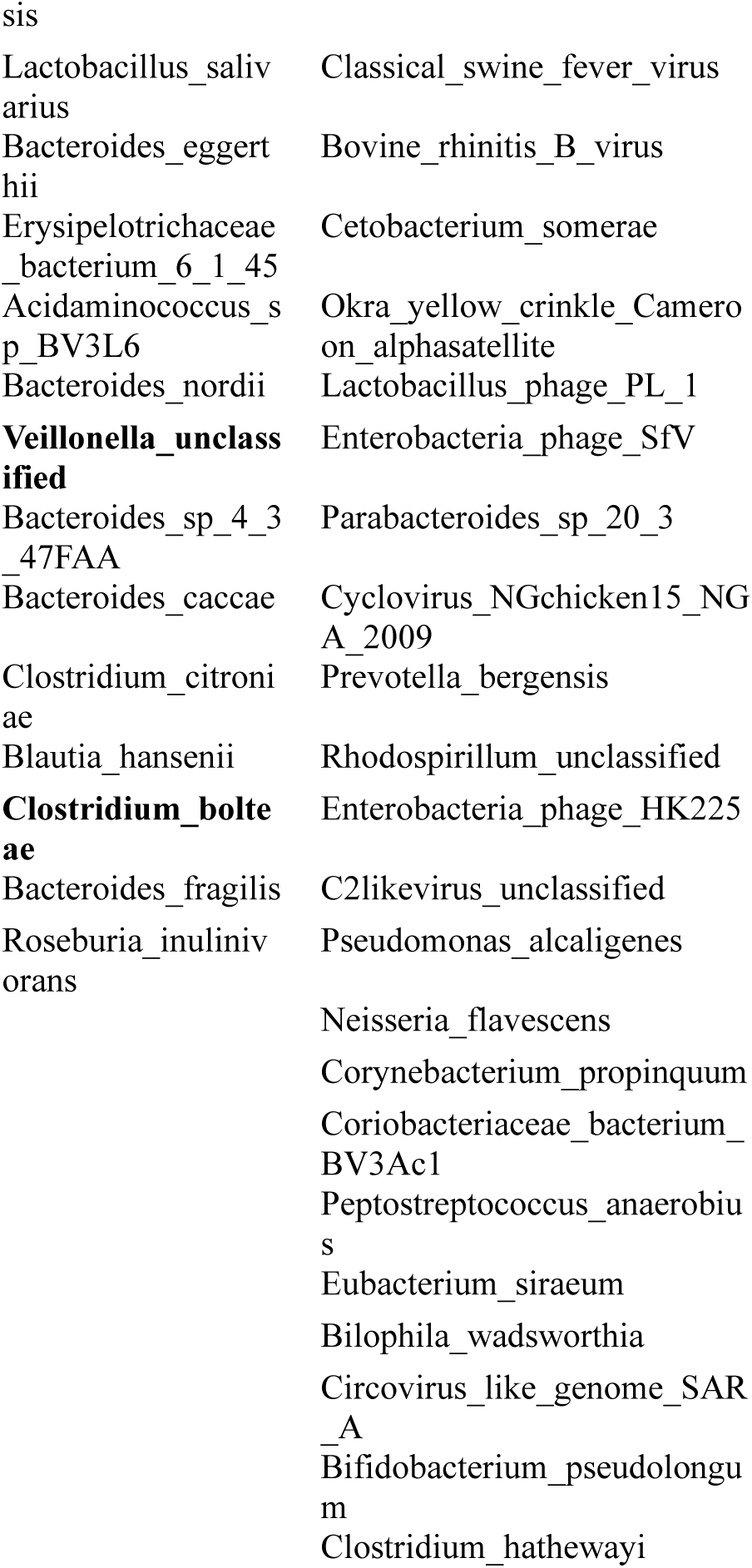
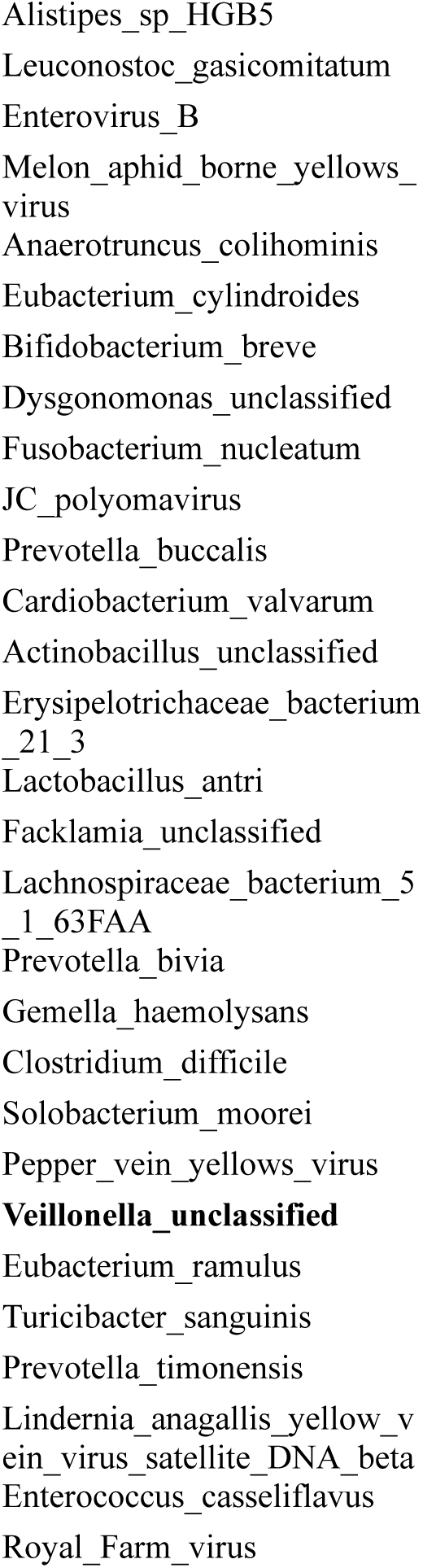
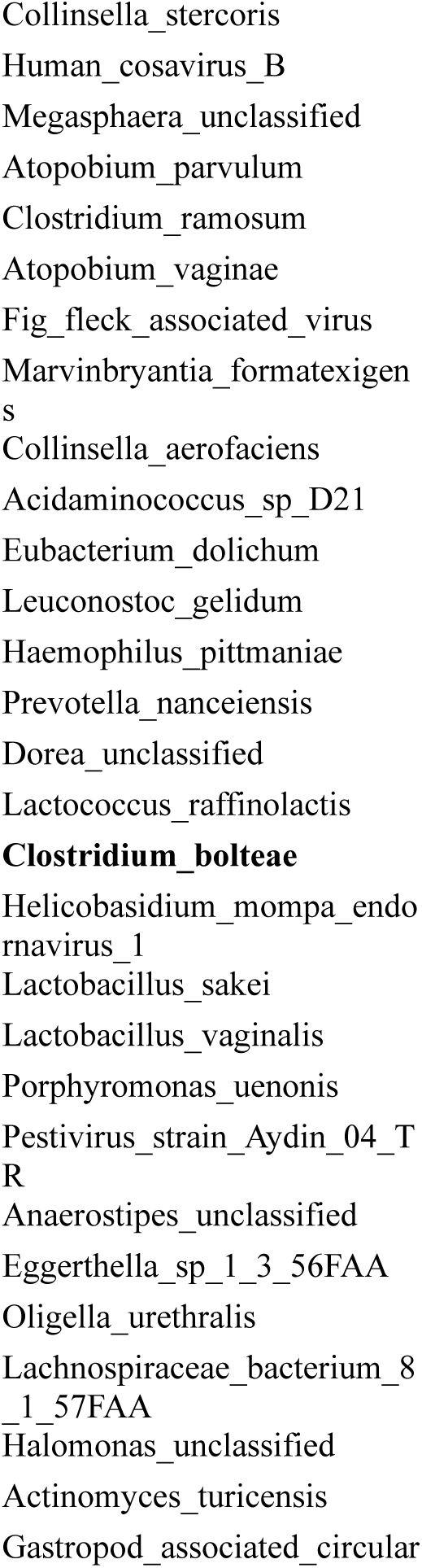
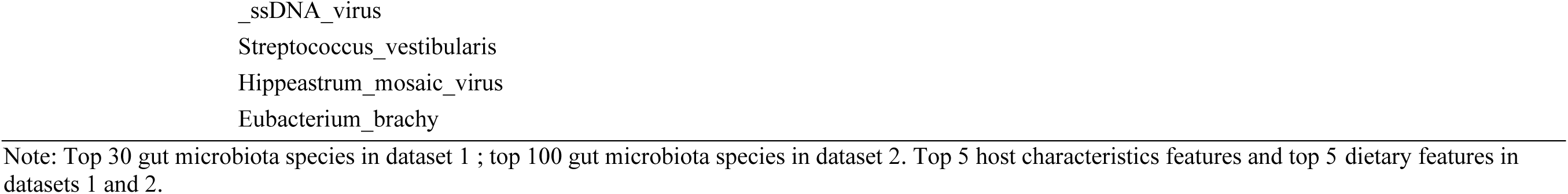
Important factors associated with blood isobutyric acids.

**Supplementary Table 11.**
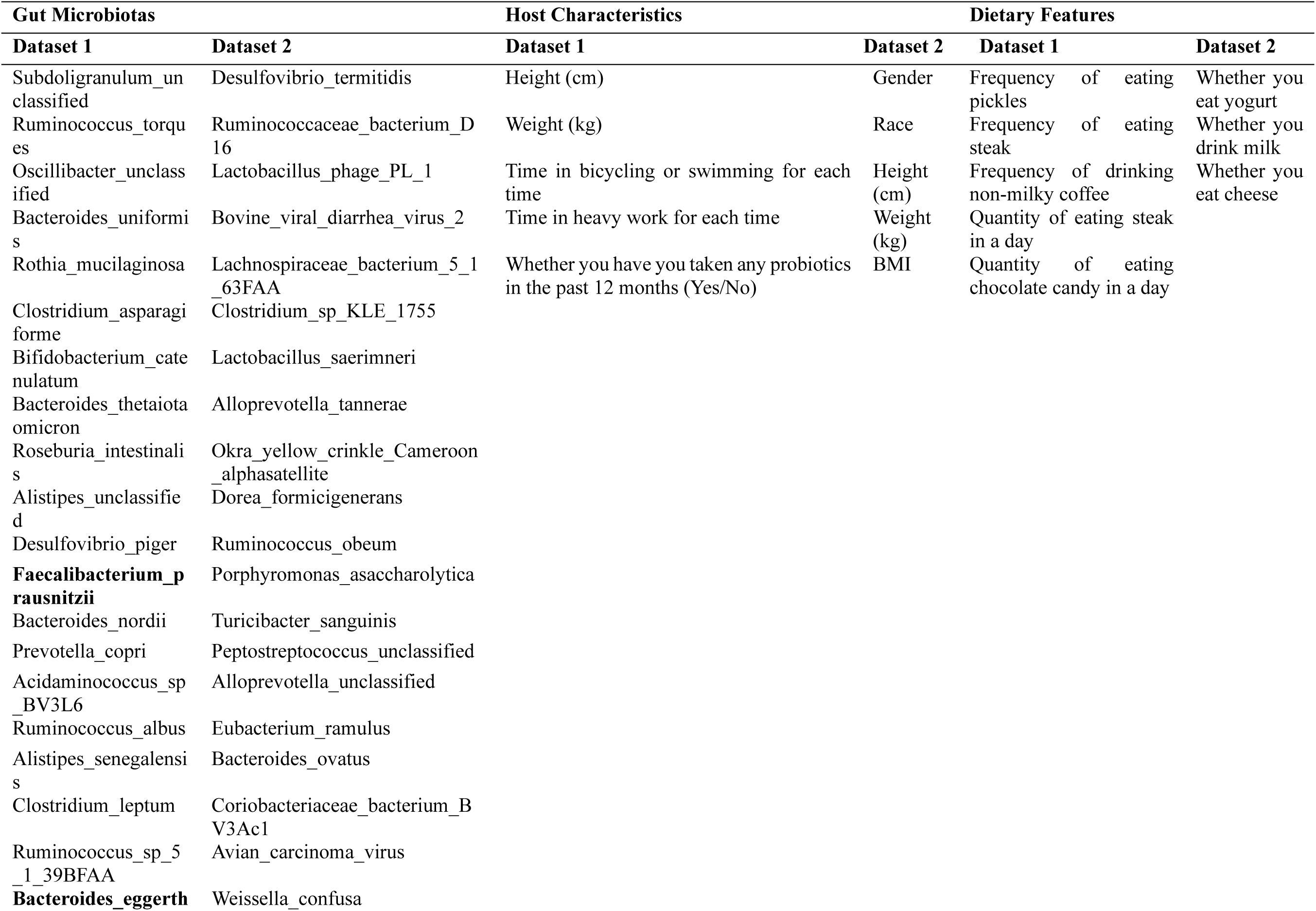
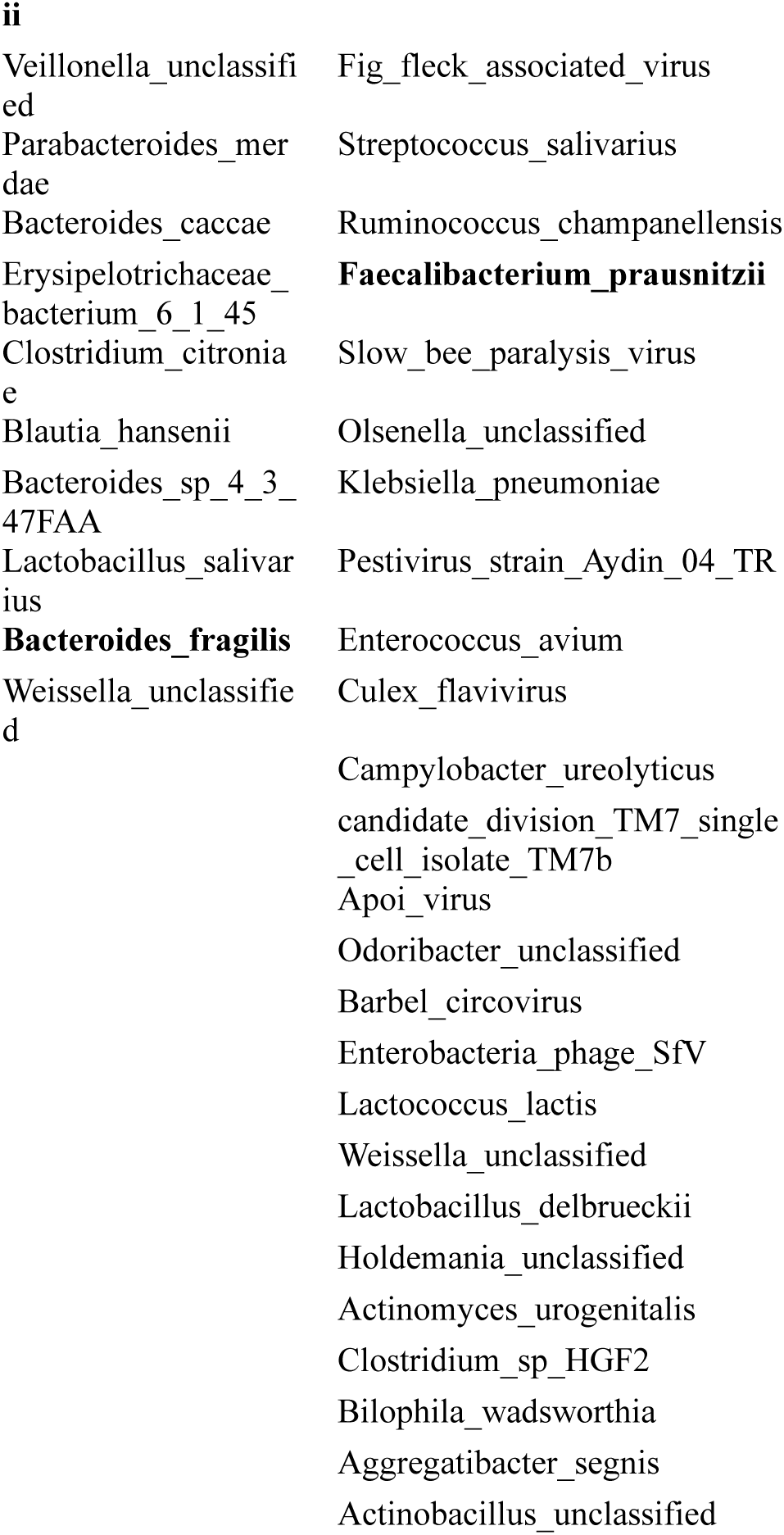
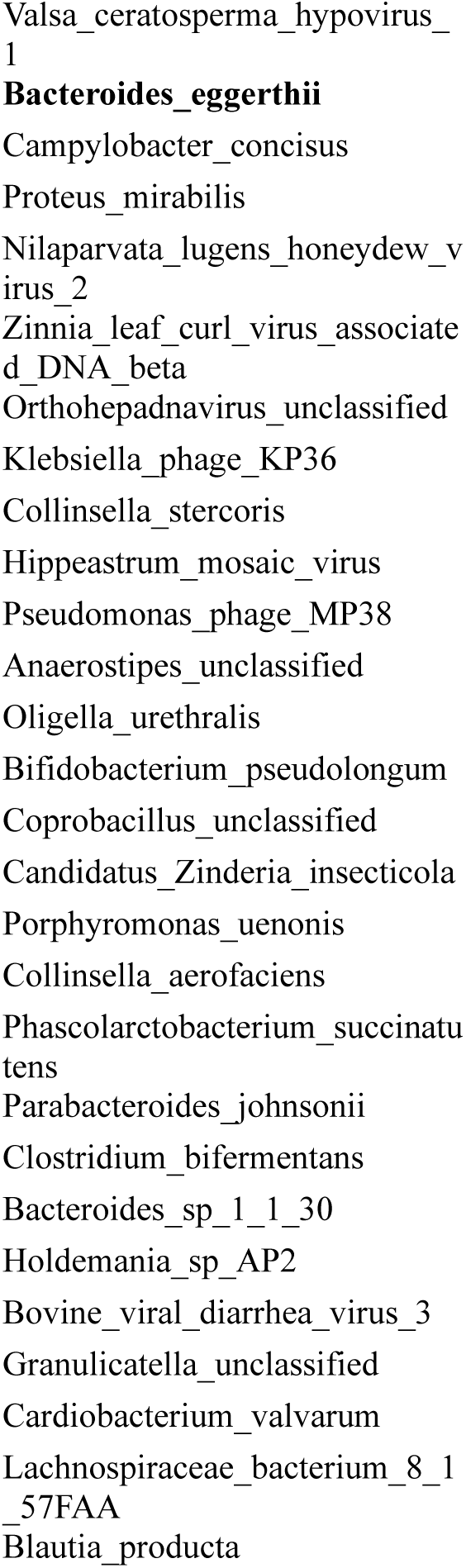
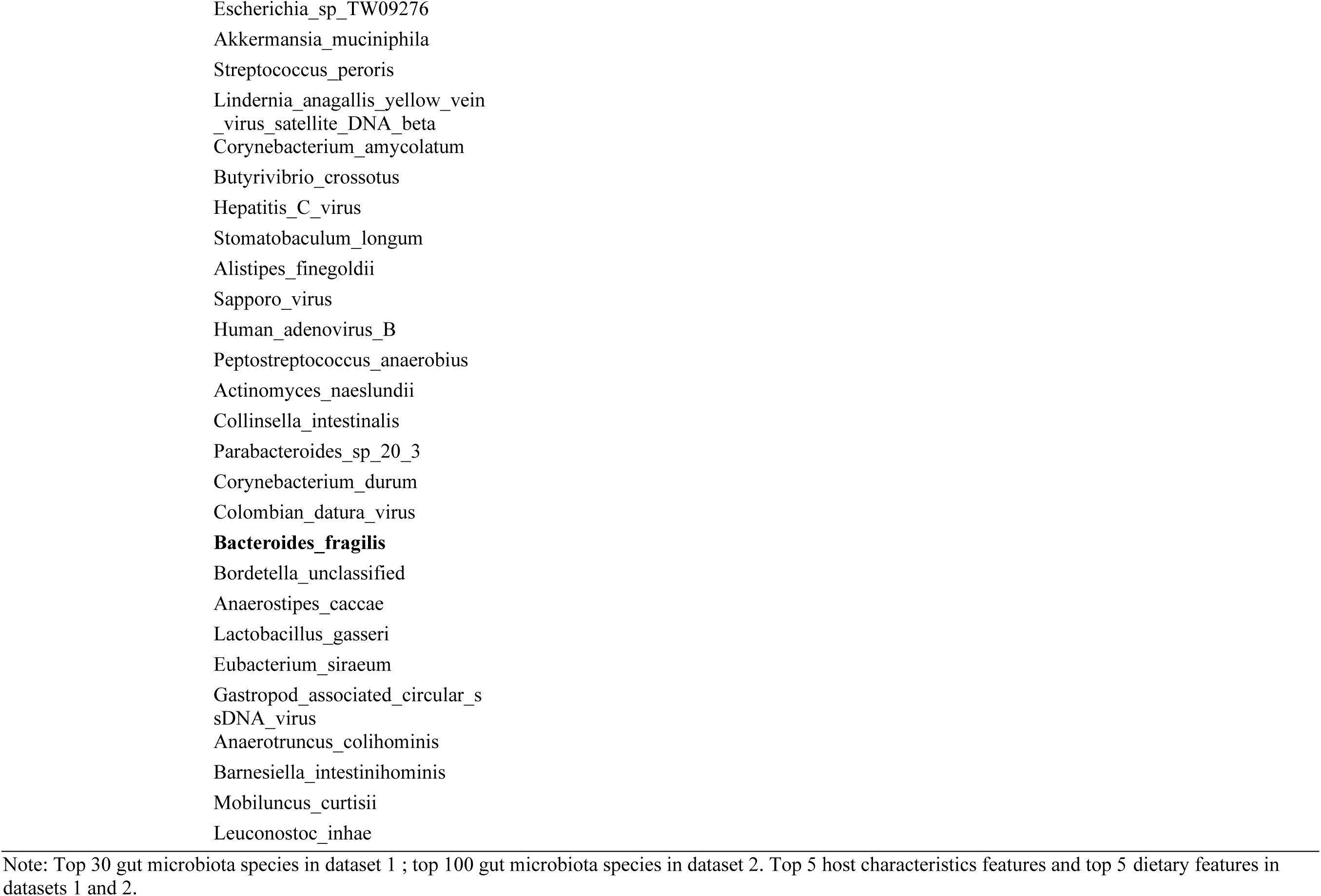
Important factors associated with blood isovaleric acids.

